# *PRDM9* losses in vertebrates are coupled to those of paralogs *ZCWPW1* and *ZCWPW2*

**DOI:** 10.1101/2021.06.08.447603

**Authors:** Maria Izabel A. Cavassim, Zachary Baker, Carla Hoge, Mikkel H. Schierup, Molly Schumer, Molly Przeworski

## Abstract

In most mammals and likely throughout vertebrates, the gene *PRDM9* specifies the locations of meiotic double strand breaks; in mice and humans at least, it also aids in their repair. For both roles, many of the molecular partners remain unknown. Here, we take a phylogenetic approach to identify genes that may be interacting with PRDM9, by leveraging the fact that *PRDM9* arose before the origin of vertebrates, but was lost many times, either partially or entirely––and with it, its role in recombination. As a first step, we characterize PRDM9 domain composition across 446 vertebrate species, inferring at least thirteen independent losses. We then use the interdigitation of *PRDM9* orthologs across vertebrates to test whether it co-evolved with any of 241 candidate genes co-expressed with PRDM9 in mice or associated with recombination phenotypes in mammals. Accounting for the phylogenetic relationship among species, we find two genes whose presence and absence is unexpectedly coincident with that of *PRDM9*: *ZCWPW1*, which was recently shown to facilitate double strand break repair, and its paralog *ZCWPW2*, as well as more tentative evidence for *TEX15* and *FBXO47*. *ZCWPW2* is expected to be recruited to sites of PRDM9 binding; its tight coevolution with *PRDM9* across vertebrates suggests that it is a key interactor, with a role either in recruiting the recombination machinery or in double strand break repair.

**Author Summary:** Our understanding of meiotic recombination in mammals has seen great progress over the past 15 years, spurred in part by the convergence of lines of evidence from molecular biology, statistical genetics and evolutionary biology. We now know that in most mammals and likely in many vertebrates, the gene *PRDM9* specifies the location of meiotic double strand breaks and that in mice and humans at least, it also aids in their repair. For both roles, however, many of the molecular partners remain unknown. To search for these, we take a phylogenetic approach, leveraging the fact that the complete *PRDM9* has been lost at least thirteen times in vertebrates and thus that its presence is interdigitated across species. By this approach, we identify two genes whose presence or absence across vertebrates is coupled to the presence or absence of *PRDM9*, *ZCWPW1* and *ZCWPW2,* as well as two genes for which the evidence is weaker, *TEX15* and *FBXO47.* ZCWPW1 was recently shown to be recruited to sites of PRDM9 binding and to aid in the repair of double strand breaks. ZCWPW2 is likely recruited to sites of PRDM9 binding as well; its tight coevolution with *PRDM9* across vertebrates suggests that it plays an important role either in double strand break formation, potentially as the missing link that recruits the recombination machinery to sites of PRDM9 binding, or in double strand break repair.

## Introduction

Meiotic recombination is initiated by the deliberate infliction of numerous double strand breaks (DSBs) in the genome, the repair of which yields crossover and non-crossover resolutions (reviewed in [1]). In mice and humans, and probably in most mammals, the localization of almost all DSBs is specified through the binding of PRDM9 [2-4]. Yet the presence of a PRDM9 binding site is far from sufficient for a DSB to be made: a number of additional factors modulate whether PRDM9 binds or act downstream of PRDM9 binding [5-7].

The mechanism by which PRDM9 directs recombination to the genome is partially understood: it binds DNA through a C2H2 zinc finger (ZF) array and contains a SET domain that tri-methylates histones H3K4 and H3K36 [8,9]. These epigenetic marks together recruit the DSB machinery, notably SPO11 (which makes the DSBs), through intermediates that remain unknown [10]. In addition to the zinc finger binding array and SET domain, most mammalian *PRDM9* genes also have two other domains, KRAB and SSXRD, whose functions are unclear.

The complete PRDM9 protein, with all four domains, originated before the diversification of vertebrates, so has been conserved for hundreds of millions of years [11,12]. Yet the entire gene has also been lost numerous times, including in birds and canids [13-15]. In these species, recombination occurs preferentially around promoter-like features, notably CpG islands [11,15–17]. A possible explanation is that in the absence of the histone marks laid down by PRDM9, the recombination machinery defaults to those residual H3K4me3 marks found in the genome, often associated with sites of transcription initiation, or perhaps simply to wherever DNA is accessible [15,18]. The same concentration of DSBs around promoter-like features is seen in *Prdm9*^-/-^ mice [18] and in a woman who carries two loss of function copies of *PRDM9* identical by descent [19]. These findings suggest that mammals that carry an intact *PRDM9* retain the mechanism to direct recombination employed by species lacking *PRDM9*, but it is normally outcompeted by PRDM9 binding.

In addition to complete losses of *PRDM9*, multiple partial losses have occurred independently (e.g., in platypus and various fish lineages), usually involving the truncation of the N-terminal KRAB and SSXRD domains [11]. Although these partial *PRDM9* orthologs evolve under selective constraint and thus must have some conserved function [11], several lines of evidence indicate that they do not direct recombination. For one, only in species with a complete *PRDM9* is the ZF unusually rapidly evolving in its binding affinity [11]. Since the rapid evolution of the ZF is thought to arise from the role of PRDM9 in recombination [3,20,21], this evolutionary pattern suggests that all four domains are required for DSB localization. Empirical data support this notion: in swordtail fish carrying one *PRDM9* ortholog that lacks KRAB and SSXRD domains as well as in a mouse model in which only the KRAB domain is knocked out, recombination events are concentrated at promoter-like features, as in species lacking *PRDM9* altogether [11,22]. Therefore, the KRAB domain at least appears to be necessary for PRDM9 to direct recombination, likely by mediating interactions with other proteins [22,23].

Conversely, the presence of a complete *PRDM9* with a rapidly evolving ZF outside of mammals [11] suggests that PRDM9 also directs recombination to the genome in these species, as has been reported for rattlesnakes [24]. Thus, at least two mechanisms for directing meiotic recombination are interdigitated within mammals as well as seemingly throughout the vertebrate phylogeny.

In addition to specifying the locations of DSBs, PRDM9 has recently been discovered to play a second role, in the downstream repair of DSBs [25-27]. In mice and humans, DSBs at which PRDM9 is bound on both homologs are more likely to be efficiently repaired and to result in a crossover; in contrast, DSBs at which PRDM9 is only bound on one of the two homologs are delayed in their repair [27,28]. If these “asymmetric” DSBs are overwhelming in number—as is the case in certain hybrid crosses in mice—this delay can lead to asynapsis and infertility [29,30].

While this second role of PRDM9 is still poorly understood, recent papers report that it is facilitated by ZCWPW1, which binds H3K4me3 and H3K36me3 [25-27] and is expressed alongside PRDM9 in single cell data from mouse testes [31]. One line of evidence that enabled the discovery of *ZCWPW1* is that although it too has been lost numerous times in vertebrates, it is found in seven clades that carry an intact *PRDM9* [26,27].

The important hint provided by the phylogenetic distribution of *ZCWPW1* points to the potential power of co-evolutionary tests to identify additional molecular partners of PRDM9. Here, we took this approach more systematically: we considered a set of 241 candidate genes that are either known to be involved in recombination in model organisms [32], associated with recombination phenotypes in a human genome-wide association study [33], or co-expressed with PRDM9 in single cell data from mouse testes [31] and tested for their co-occurrence with *PRDM9* across 189 vertebrate species. After verifying our initial gene status calls in whole genome data and, for a subset of species, RNA-seq data, we identified the paralog of *ZCWPW1*, *ZCWPW2*, as co-evolving with *PRDM9* and found more tentative evidence for two additional genes, *TEX15* and *FBXO47*.

## Results

### A revised phylogeny of PRDM9

We previously reported that the complete *PRDM9* gene, including KRAB, SSXRD and SET domains, arose before the origin of vertebrates and was lost independently a number of times, both in its entirety and partially (through the loss of its N-terminal domains; [11]). Here, we leverage the independent losses of *PRDM9* in order to identify genes that are co-evolving with *PRDM9*––specifically, that tend to be present in the same species as *PRDM9* and lost (partially or entirely) when *PRDM9* is no longer complete.

As a first step, we characterized the phylogenetic distribution of *PRDM9* in light of new genome sequences published since our initial analysis [11]. To this end, we created a curated dataset of 747 vertebrate *PRDM9* sequences by analyzing publicly available protein sequences from Refseq [34], whole genome sequences, and RNA-seq data from testes samples, as well as four RNA-seq datasets from testes samples that we generated (see Methods, **Figure S1**, **Tables S1-3**). For this analysis, we defined *PRDM9* orthologs as complete if they contain both KRAB and SET domains; we did not consider the SSXRD domain, because its short length makes its detection at a given e-value threshold unreliable, or the ZF array, because its repetitive structure makes it difficult to sequence and assemble reliably.

Across 446 species, we identified 221 species with at least one complete *PRDM9* ortholog and 225 species without a complete *PRDM9* ortholog (**Figure 1**, **Table S4**). Notably, we were able to uncover complete *PRDM9* orthologs in a number of species for which we had previously predicted partial or complete losses [11], including in the Tasmanian devil (*Sarcophilus harrisii*), the atlantic cod (*Gadus morhua*), and the atlantic herring (*Clupea harengus*), as well as in a handful of placental mammals that we had previously only investigated using RefSeq (see **Table S4** for details). We also found a complete *PRDM9* ortholog in caecilians and in two species of frogs, suggesting that the previously reported loss of *PRDM9* in amphibians [11] reflects at least one loss in salamanders and more than one independent loss in frogs. We note, finally, that by the approach taken here, the *PRDM9* ortholog from the Australian ghostshark (*Callorhinchus milii*) is considered to be complete (in contrast to in Baker et al. 2017, where we also relied on the SSXRD domain; see **Table S4** for details).

**Figure 1.**
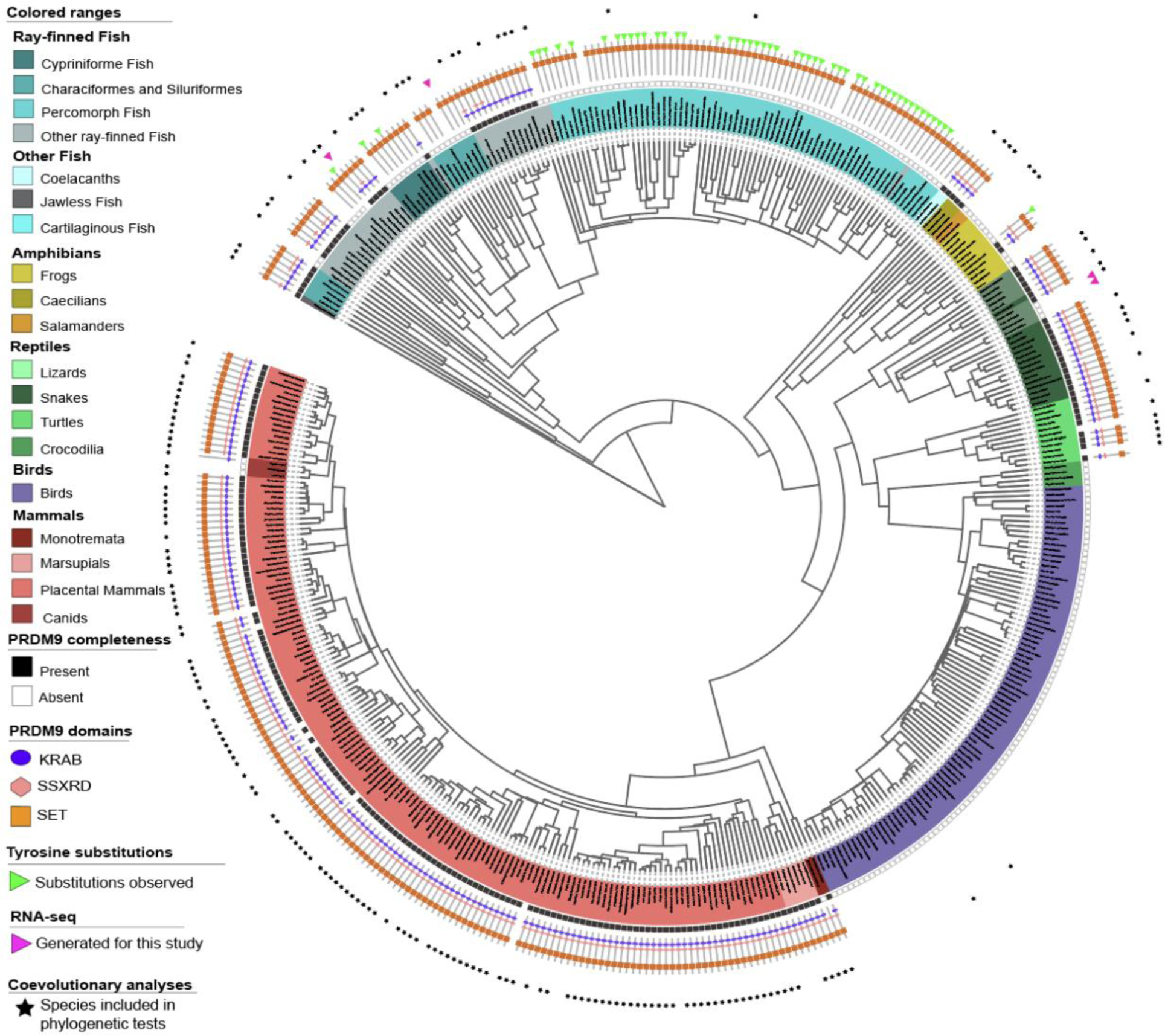
The phylogenetic distribution of PRDM9 and its domain architecture across vertebrates. The inferred *PRDM9* status of 432 vertebrate species. Branch lengths were computed based on the TimeTree database. For 28 species not present in the database, we used branch length information from a close evolutionary relative; for 14 species in which we made PRDM9 calls, we were unable to find such a substitute, so they are not represented. Different vertebrate clades are indicated by colored segments, with salmon for mammals, cyan for fish, mustard for amphibians, green for reptiles, and purple for birds. In the inner circle, squares indicate whether *PRDM9* is complete (filled black) or incomplete/absent (empty black); for species with an uncertain *PRDM9* status, no box is shown. The PRDM9 domain architecture of each species is shown with a cartoon, where the presence of a KRAB domain is indicated in blue, of SSXRD in pink, and of the SET domain in orange. Green triangles indicate species that only carry *PRDM9* orthologs with substitutions at putatively important catalytic residues in the SET domain (see **Table S4**). The tree was drawn using itool (https://itol.embl.de/); an interactive version is available at https://itol.embl.de/shared/izabelcavassim.

Given the phylogenetic relationships among species given by the TimeTree tool (http://timetree.org/; [35]), we inferred 23 putative complete or partial losses of *PRDM9* across the 446 vertebrates considered (**Figure 1**, **Table S4**). These putative losses include six previously reported ones [11,13–15], each observed in two or more closely related species: in percomorph fish, cypriniformes fish, characiformes and siluriformes fish, osteoglossomorpha fish, birds and crocodiles, and canids. In addition, independent work supports our finding of a partial loss of *PRDM9* in the platypus (*Ornithorhynchus anatinus*) (J. Hussin and P. Donnelly, personal communication). In turn, the putative losses of PRDM9 in polypteriformes fish, salamanders, and in three clades of frog species (*Xenopus, Dicroglossidae* and *Bufonidae*) were each supported by the absence of PRDM9 in the genomes of two or more closely related species. We were further able to verify the absence of PRDM9 in two *Xenopus* frogs and in two salamanders using RNA-seq data from testes: despite sufficient power to detect a set of six highly conserved meiotic genes in each species, we did not detect the expression of any complete PRDM9 orthologs (**Table S3**).

We also failed to find *PRDM9* in RefSeq or the whole genome sequence of the green anole (*Anolis carolinensis*). We verified this absence of *PRDM9* by collecting RNA-seq data from testes in the green anole as well as in the fence lizard (*Sceloporus undulatus*), for which neither a Refseq nor a genome sequence were available at the time. Despite sufficient power to detect a set of six highly conserved meiotic genes, we were unable to detect PRDM9 expression in either species (**Figure S2-3**, **Table S3**). Given the presence of a complete *PRDM9* in bearded dragons (*Pogona vitticeps*), it appears that this loss of *PRDM9* occurred in a lineage basal to the common ancestor of green anoles and fence lizards, over 99 Mya but less than 157 Mya (**Figure S5**).

The remaining 10 putative absences of *PRDM9* are observed in single species; we were unable to verify the calls using testis RNA-seq data, so their *PRDM9* status remains uncertain. Thus, in total, we identified at least 13 independent *PRDM9* losses in vertebrates, and possibly as many as 23 (**Figure 1**, **Table S4**). The 13 losses are all relatively old (**Figure S4**): the most recent case manifest in these data is either the one that happened in the branch leading to platypus or the one in canids, which could be as recent as 14.2 Mya (**Figure S4**).

### Identifying genes co-evolving with PRDM9

We selected 193 candidate genes based on their co-expression with PRDM9 in single cell RNA-seq data from mouse testes (specifically, in component 5; see Methods [31]) (**Figure S6A-B**). To this set, we added any gene associated with variation in recombination phenotypes in humans [33] as well as genes known to have a role in mammalian meiotic recombination from functional studies (summarized in [32]). Together, these three sources provided a total of 241 genes to evaluate for possible co-evolution with *PRDM9* (**Table S5**, **Figure S6C**).

We evaluated the presence or absence of these 241 genes across the NCBI RefSeq database of 189 species. These 189 species were downsampled from the larger phylogenetic tree to preserve at most three species with high quality genomes below each *PRDM9* loss, thereby minimizing phylogenetic signals of genome quality. The phylogeny includes representative species for 11 of the 13 inferred PRDM9 losses (see Methods, **Figure S7**). Species of *Bufonidae* frogs and salamanders were not included due to the absence of available gene annotations; moreover, due to the lack of gene annotations for frog species with PRDM9, within these 189 species, the losses in *Xenopus* and *Dicroglossidae* frogs cannot be distinguished from a single event.

We encoded a gene as present when it contained all the domains found in four representative vertebrates with a complete *PRDM9* and absent if it lacked one or more of those domains (see Methods). Many of the 241 genes are present in every sampled vertebrate and hence provide no information in our co-evolutionary test of presence and absence: specifically, we found apparently complete orthologs for 102 candidate genes in all 189 species used in the phylogenetic test. We therefore focused on the remaining 139 genes, each of which has been lost at least once among vertebrate species evaluated here; the matrix of 189x139 gene status calls is presented in **Table S6**.

We tested for the co-evolution of *PRDM9* and each candidate gene by comparing a null model with independent rates of gains and losses of *PRDM9* and of the focal gene to an alternative model in which the state transition rates of the two genes are dependent on one another, using the maximum likelihood approach within *BayestraitsV3* [36,37] (**Table S7**, **Figure S8**). By this approach, we identified nine significant hits at the 5% level (uncorrected for multiple tests): in order of increasing p-values, *ZCWPW1, MEI1*, *ZCWPW2*, *TEX15*, *FBXO47*, *ANKRD31*, *NFKBIL1*, *SYCE1*, and *FMR1NB*. We focused on the top five, for which the false discovery (FDR) value is below 50% (**Table 1**, **Figure 2A**).

**Figure 2.**
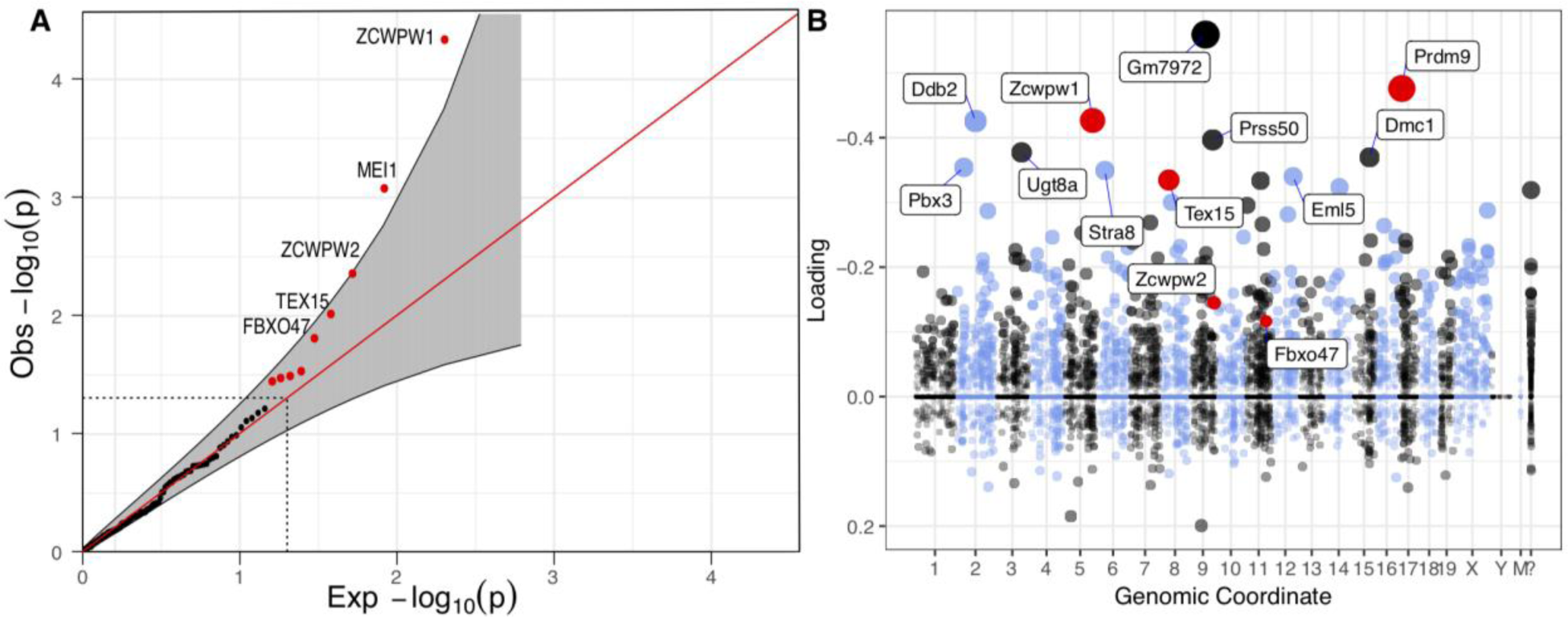
Phylogenetic tests and genes co-expressed with PRDM9 in single cell mouse testes data. **(A)** Quantile-Quantile plot of the p-values obtained from the phylogenetic tests run on 139 genes that appeared to have been lost at least once in the 189 vertebrate species considered. Genes that are significant at the 5% level are shown in red (outside the dashed lines) and a pointwise 95% confidence interval is shown in grey. Genes with a FDR ≤ 50% are annotated. **(B)** Loadings for one of 46 components (component 5) inferred from single cell expression data in mouse testes [31], in which PRDM9 is most highly expressed. The dot sizes are proportional to the square of the absolute value of the loading. *PRDM9* and the three genes identified in our phylogenetic tests with p<0.05 are shown in red. Mouse genomic coordinates are displayed. Panel B was made from summary statistics provided by [31], using SDAtools (https://github.com/marchinilab/SDAtools/).

**Table 1.** Results of phylogenetic tests. We focused on the five genes that had a false discovery rate (FDR) ≤ 50%, improved the ortholog status calls, and reran the phylogenetic tests for four of them (all but *MEI1*, which turned out to be present in all species considered; see Methods). Gene source refers to the criterion by which the gene was originally included among our lists of candidates: (1) It is co-expressed with PRDM9 in single cell mouse testes data [31] or (2) variants assigned to the gene are associated with variation in recombination phenotypes in humans [33] or (3) the gene was previously known to have a role in mammalian meiotic recombination from functional studies [32] (see Methods).

| Gene | Position<br>(human<br>coordina<br>te) | Gene<br>source | LogLik<br>H0 | LogLik<br>Ha | P-value | FDR | P-value for<br>improved<br>status calls |
| --- | --- | --- | --- | --- | --- | --- | --- |
| *ZCWPW1* | chr7:<br>1<br>00400826 | 1 | -65.941 | -53.35647 | 4.651e-05 | 0.0064 | 1.948e-03 |
| *MEI1* | chr22:<br>4<br>1699503 | 3,1 | -54.200 | -44.779 | 8.442e-04 | 0.0586 | NA |
| *ZCWPW2* | chr3:<br>2834872<br>1 | 1 | -67.711 | -60.146 | 4.437e-03 | 0.2055 | 5.171e-06 |
| *TEX15* | chr8:<br>3<br>0831544 | 1 | -138.430 | -131.764 | 9.760e-03 | 0.3391 | 8.682e-02 |
| *FBXO47* | chr17:<br>3893643<br>2 | 2 | -53.678 | -47.559 | 1.566e-02 | 0.4354 | 1.566e-02 |

We sought to verify the phylogenetic distribution of the top genes by developing curated datasets of high confidence orthologs, as we had for *PRDM9* (see Methods; **Figure 3**, **Tables S1** and **S8, Figure S8**). In doing so, we were able to identify *MEI1* orthologs from the whole genome assemblies of each species missing *MEI1* in our initial dataset, resulting in the presence of *MEI1* in every species considered (**Table S1**); thus, it appears that its inferred co-evolution with *PRDM9* based on Refseq calls is artifactual (see Methods). Rerunning the phylogenetic test on the curated ortholog sets for the remaining four genes, *TEX15* is no longer significant at the 5% level (*p*=0.086), possibly because the curation uncovered an intact *TEX15* ortholog in anoles. *ZCWPW1* and *ZCWPW2* are still highly significant; for *FBXO47,* the curation did not reveal any discrepancy with the initial calls, so the p-value remains the same.

**Figure 3.**
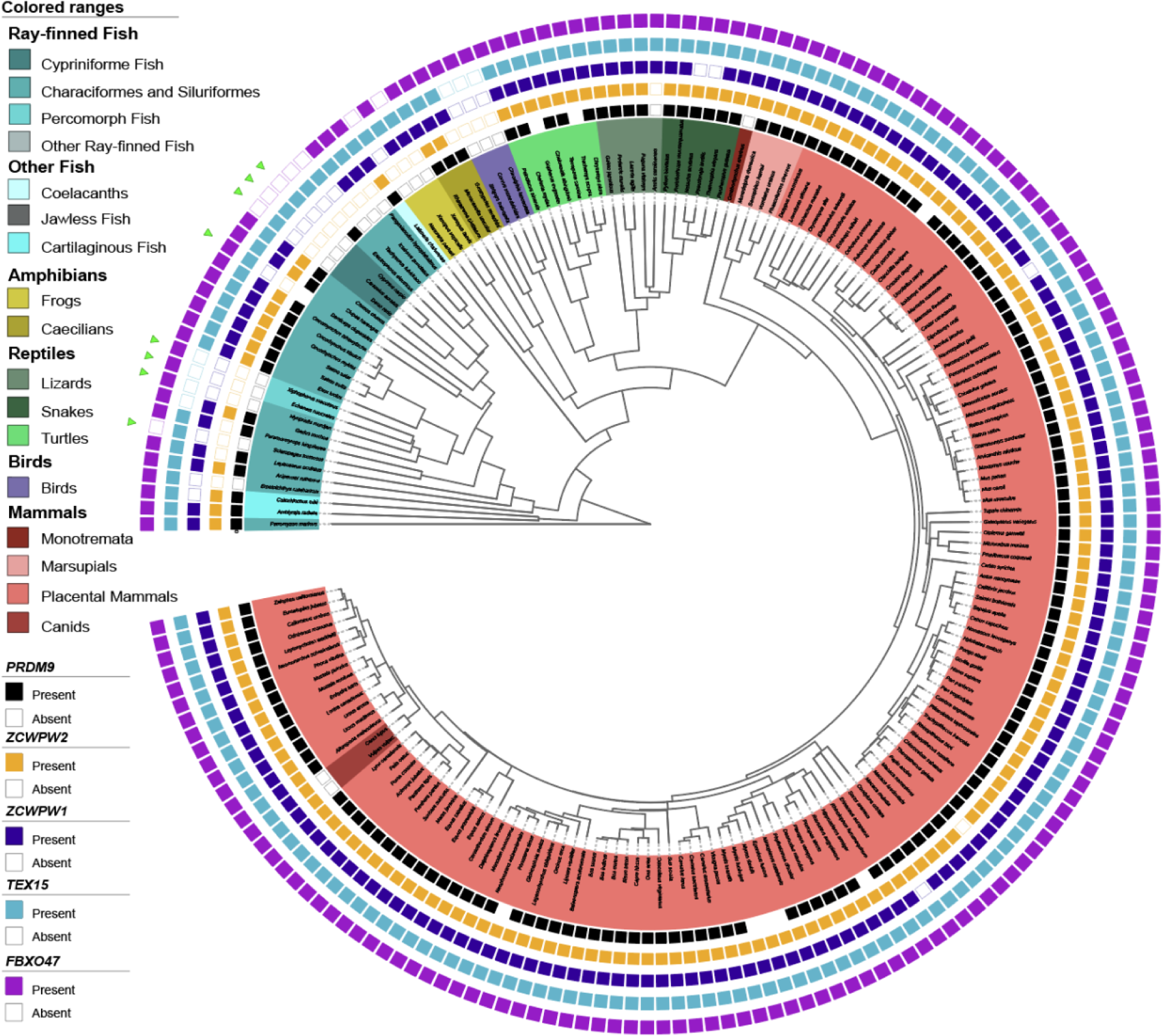
The phylogenetic distribution of *PRDM9* and co-evolving genes across 189 species. Filled teal and empty teal squares indicate whether *PRDM9* is present or absent, respectively (see Methods). If nothing is indicated, the status of *PRDM9* is uncertain. Likewise, filled orange and empty squares indicate whether *ZCWPW2* is present or absent/incomplete; filled and empty navy squares indicate whether *ZCWPW1* is present or absent/incomplete; filled and empty light blue squares indicate whether *TEX15* is present or absent/incomplete; and filled and empty light purple squares indicate whether *FBXO47* is present or absent/incomplete. Green triangles indicate species that only carry *PRDM9* orthologs with substitutions at putatively important catalytic residues in the SET domain (see **Table S4**). The status of candidate genes (for which FDR ≤ 50%; Figure 2A) was re-evaluated based on a search of gene models within whole genome sequences (see Methods); updated p-values for the phylogenetic test are shown in Table 1. The tree was drawn using itool (https://itol.embl.de/); an interactive version is available at https://itol.embl.de/shared/izabelcavassim.

Our approach therefore uncovered two genes with clear-cut evidence of co-evolution with *PRDM9*, the paralogs *ZCWPW1* and *ZCWPW2*, and more tentative support for two others, *TEX15* and *FBX047*. *ZCWPW1*, *ZCWPW2* and *TEX15* were among our initial list of 241 candidate genes because they are co-expressed with PRDM9 in single cell testis data from mouse [31] (**Figure 2B**; **Figure S6**). *FBX047* was not included by that criterion but because missense mutations in the gene are associated with recombination rate variation in the total genetic map length in humans, in both males and females [33]. Nonetheless, the expression of *FBXO47* in mice is testis-specific [38], and the gene is expressed in the component in which *PRDM9* had the highest loading (albeit with a smaller loading [31] **Figure 2B**; see also [39]).

Like *PRDM9*, *ZCWPW1*, *ZCWPW2, FBXO47* and *TEX15* are inferred to have been present in the common ancestor of vertebrates. Below we describe the distribution of each of the four genes across the phylogeny of 189 species and the patterns that give rise to the evidence of statistical association with *PRDM9*--in particular, the correspondence between their distributions and that of 11 well supported losses of *PRDM9*, as well as of 9 species for which the status of *PRDM9* is uncertain.

### *ZCWPW1* and *PRDM9* co-evolution

Our finding that *ZCWPW1* is co-evolving with *PRDM9* (*p*=0.0019 in the curated set; Table 1) is in line with previous reports of an association in vertebrates between the presence and absence of *ZCWPW1* and *PRDM9* orthologs [26,27]. Here, we found an even tighter coupling of *PRDM9* and *ZCWPW1* than previously documented. Specifically, we inferred 12 losses of *ZCWPW1* among 189 species used in our phylogenetic test, distributed across 17 species that lack *ZCWPW1* entirely and two species carrying partial *ZCWPW1* genes (with the PWWP domain but not the zf-CW domain; **Table S8**).

Seven of the *ZCWPW1* losses occur among the 11 well supported losses of *PRDM9*: in cypriniformes fish, percomorph fish (*Euacanthomorphacea*), siluriformes fish, polypteriformes fish, osteoglossomorpha fish, birds, and *Dicroglossidae* frogs. An additional *ZCWPW1* loss occurred in the denticle herring (*Denticeps clupeoides*), a species for which the status of *PRDM9* is uncertain. The remaining four losses of *ZCWPW1* seem to break the pattern, in that they occur in lineages containing a complete *PRDM9* gene. However, three are observed only in a single species and may be spurious. Therefore, across the tree, there is only one well supported case of a taxon with an intact *PRDM9* that has nonetheless lost *ZCWPW1*, supported by two closely related species, the tiger snake (*Notechis scutalus*) and the eastern brown snake (*Pseudonaja textilis*) (see **Table S8** for details).

In mice as well as human cell lines, ZCWPW1 binds two marks laid down by PRDM9: the zf-CW domain binds H3K4me3 and the PWWP domain H3K36me3 [40-42]. Thus, the co-evolution across vertebrates likely reflects a conserved molecular interaction between ZCWPW1 and PRDM9 as reader and writer of these dual histone modifications.

### *ZCWPW2* also co-evolves with *PRDM9*

Intriguingly, the strongest association with the presence or absence of *PRDM9* is that of the paralog of *ZCWPW1*, *ZCWPW2* (*p*=5x10^-6^; **Table 1**). Among the 189 species, there are 12 independent losses, distributed across 21 species that appear to lack *ZCWPW2* altogether and three that contain partial *ZCWPW2* genes (two with the PWWP domain but not the zf-CW domain, and one with the reverse; **Table S8**).

Six of the *ZCWPW2* losses occur among the 11 well supported losses of an intact PRDM9: in percomorph fish, polypteriformes fish, *Xenopus* frogs, *Dicroglossidae* frogs, birds, and the green anole. In order to distinguish whether the absence of *ZCWPW2* in *Xenopus* and *Dicroglossidae* frogs reflects a single loss or multiple events, we investigated the status of *ZCWPW2* in an additional species of frog with *PRDM9* (*Ranitomeya imitator*). We were able to successfully identify a complete *ZCWPW2* ortholog in this species, suggesting that *ZCWPW2* has indeed been lost at least twice within frogs, possibly coincidentally with *PRDM9* in each case. *ZCWPW2* is also absent in a clade encompassing cypriniformes fish and siluriformes fish, as well as the electric eel (*Electrophorus electricus*), which has an intact *PRDM9*. This phylogenetic distribution suggests that the loss of *ZCWPW2* may have occurred before the losses of *PRDM9* in both cypriniformes fish and siluriformes fish. Also suggestive of this order of loss, *ZCWPW2* is absent in osteoglossomorpha fish (the Asian arowana, *Scleropages formosus*); in this case, the gene is also absent from the closest evolutionary relative in the tree, the elephantfish (*Paramormyrops kingsleyae*), which carries *PRDM9*.

Among the nine species for which the status of *PRDM9* is uncertain, *ZCWPW2* is absent in the denticle herring (*Denticeps clupeoides*). The remaining three cases of *ZCWPW2* loss are each observed in a single species carrying an intact PRDM9, without supporting lines of evidence. In summary, in the few cases with *PRDM9* but not *ZCWPW2*, we cannot verify the loss of *ZCWPW2*; conversely, the only species with *ZCWPW2* but that clearly lack *PRDM9* are canids and the platypus, the two lineages that experienced the most recent losses of *PRDM9* (see **Table S8** for details).

Like its paralog, ZCWPW2 contains zf-CW and PWWP domains, predicted to bind H3K4me3 and H3K36me3, respectively (**Figure 4A**, **Figure S10**). As in ZCWPW1 ([27], [26]), these domains are highly conserved, especially at residues with predicted binding properties (**Figure 4B-C**), suggesting that ZCWPW2 is also recruited to sites of PRDM9 binding.

**Figure 4.**
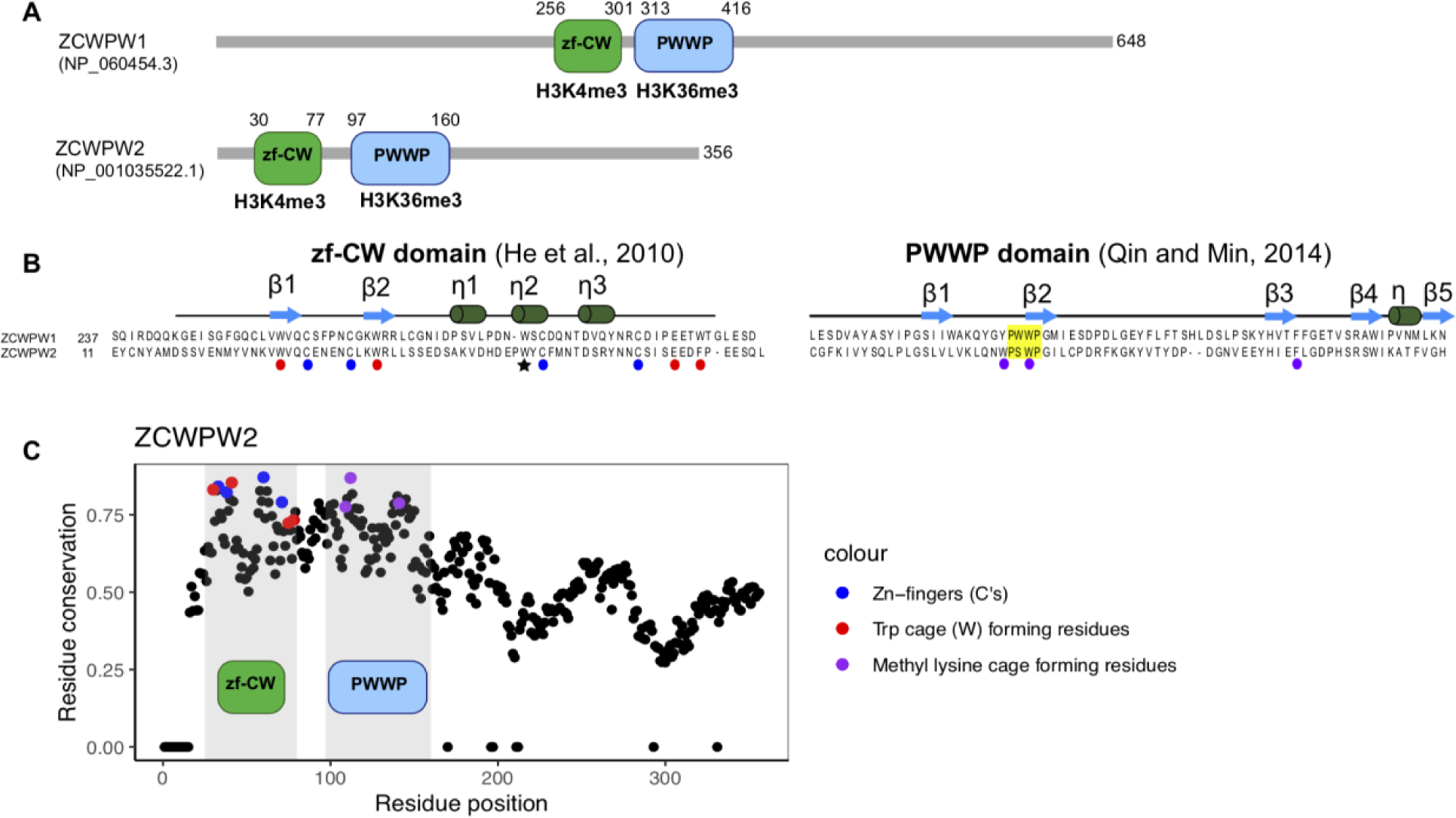
Domain architecture and conservation of *ZCWPW1* and *ZCWPW2*. **(A)** Amino acid sequence and domain structure composition of genes *ZCWPW1* and *ZCWPW2* in humans. **(B)** The ZF-CW domain structure includes the fingers (residues indicated by blue circles) and an aromatic cage (red) expected to bind to H3K4me3 [73], and the star indicates the third Trp residue that is thought to stabilize the fold by hydrophobic interactions [73]. The PWWP domain (yellow) is expected to bind to histone H3K36me3 through a hydrophobic cavity composed of three aromatic residues (purple) [74]. The secondary structures of zf-CW and PWWP domains are represented above sequences. **(C)** Conservation of residues in *ZCWPW2* across vertebrates, with those residues recognizing modifications on the histone tail colored in blue, red and purple.Positions in the *ZCWPW2* alignment with >30% of gaps were ignored and the conservation score was set to 0.

### The distribution of *FBXO47* and *TEX15* orthologs

We identified two additional genes, *FBX047* and *TEX15*, that may be co-evolving with *PRDM9*: using the curated calls, *p=*0.016 and p=0.087, respectively (**Table 1**). TEX15 is co-expressed with PRDM9 in two components inferred from single cell data from mice, active during pre-leptotene and zygotene (**Figure S6**). The statistical evidence for co-evolution stems from the fact that *TEX15* is missing in two taxa lacking *PRDM9*: birds and percomorph fish. *TEX15* is also absent in the Atlantic cod (*Gadus morhua*), suggesting that the loss of *TEX15* that led to its absence in percomorph fish occurred before that of *PRDM9*. All of the other 189 species considered have an intact *TEX15* (see **Table S8** for details).

The statistical evidence is a bit stronger for *FBX047*, which has been lost five times in the absence of *PRDM9*: in cypriniformes fish, osteoglossomorpha fish, siluriformes fish, and in *Xenopus* and *Bufonidae* frogs. Intriguingly, *FBXO47* is additionally absent in the electric eel, a species that carries a complete *PRDM9* gene, but lacks both *ZCWPW1* and *ZCWPW2*. Testing for the co-evolution of the candidate genes with each other, a null model in which the state transitions of *FBXO47, ZCWPW1* and *ZCWPW2* are independent is rejected for all pairs of genes (maximal p<6x10^-3^; **Table S9A**), and p-values are lower for *FBXO47* and *ZCWPW1*, or *FBXO47* and *ZCWPW2*, than for *FBXO47* and *PRDM9*.

In summary, by extending the reconstruction of *PRDM9* to 446 vertebrate species, we identified thirteen losses that are supported by more than one species or by independent evidence, and possibly as many as 23. Focusing on a subset of 189 species that capture eleven state transitions of *PRDM9*, we tested whether *PRDM9* transitions coincide with those of 139 candidate genes lost at least once across vertebrates. After carefully vetting the ortholog calls for our top five signals, we identified two genes that are clearly co-evolving in their presence and absence with *PRDM9*, *ZCWPW1* and its paralog *ZCWPW2,* and two for which the evidence is weaker: *FBXO47* and most tentatively, *TEX15*.

## Discussion

### Dual roles of PRDM9 across vertebrates

We had previously hypothesized that PRDM9 plays a role in directing recombination not only in mammals but across vertebrates, based on the presence of an intact ortholog across vertebrates with a rapidly-evolving zinc finger [11]. Consistent with our prediction, there is tentative evidence for the influence of PRDM9 binding on recombination in rattlesnakes [24]. That a gene with a known role in recombination, *ZCWPW1* co-evolves with *PRDM9* across vertebrates lends further support to this hypothesis.

The precise nature of the molecular interactions between PRDM9 and ZCWPW1 remains unknown, but recent evidence suggests that ZCWPW1 interacts with PRDM9 to facilitate the repair of PRDM9-dependent DSBs: notably, *Zcwpw1*-/- male mice and older female mice are sterile [27,43] and exhibit defects in their ability to repair DSBs [25-27]. In turn, the genomic locations of DSBs are not altered in *Zcwpw1*-/- mice, indicating that the gene does not play a role in DSB positioning [25-27]. In light of these experimental results, the co-evolution of *PRDM9* with *ZCWPW1* across vertebrates indicates that PRDM9 likely plays a role in the efficient repair of DSBs not only in mice and humans [25,26,44,45], but across the vertebrate phylogeny.

### Nature of the co-evolution of candidate genes with PRDM9

If a gene interacts with PRDM9 by reading its histone modifications, as is the case for ZCWPW1 [25-27] and likely ZCWPW2 (**Figure 4**), and has no other roles, we would expect that gene to be dispensable in species that no longer have an active PRDM9 SET domain. Previous papers reported that *ZCWPW1* is more likely to be missing from ray-finned fish with substitutions in catalytic tyrosine residues of the SET domain, in addition to clades lacking the entire *PRDM9* gene [26,27]. In our analysis, we find that both *ZCWPW1* and *ZCWPW2* are more likely to be absent from species carrying only *PRDM9* orthologs with substitutions in at least one catalytic tyrosine residue, as well as those lacking *PRDM9* altogether (**Figure 3**).

While this pattern suggests a dependence of *ZCWPW1* and *ZCWPW2* on the intact catalytic activity of PRDM9, the interpretation is complicated by the fact that all species with substitutions at the tyrosine residues in all *PRDM9* copies are also carrying only partial *PRDM9* orthologs lacking KRAB and SSXRD domains, and nearly all species with conserved tyrosine residues also carry a complete copy of *PRDM9*. In that regard, the few exceptions are informative: among species with confident *PRDM9* calls, the platypus and siluriformes fish carry *PRDM9* orthologs putatively missing the KRAB domain but with intact tyrosine residues. *ZCWPW2* is absent from all three considered siluriformes fish species while *ZCWPW1* is absent from one. Thus, the presence of *ZCWPW1* and *ZCWPW2* may depend on that of the KRAB domain rather than, or in addition to, the tyrosine residues remaining intact.

Similar considerations suggest that in the rare lineages where *ZCWPW1*, *ZCWPW2, FBX047* and *TEX15* are present in the absence of *PRDM9*, we might expect the genes to be under relaxed selective constraint. To examine this prediction, we tested whether *ω* = *dn*/*ds* was higher in lineages without a complete *PRDM9* (where dn is the rate of non-synonymous substitutions and ds the rate of synonymous substitutions; see Methods). For *ZCWPW1*, there was no evidence for a relaxation of selection (*p*>0.13; **Table S10**). The intriguing exception is in platypus, one of the few species that has a SET domain with intact tyrosine residues but is lacking the KRAB domain (*p*=0.038; in all other cases, *p*>0.13; **Table S10**). This observation lends further support to the notion that the conservation of *ZCWPW1* may depend on the KRAB domain rather than, or in addition to, the tyrosine residues of *PRDM9*.

For *ZCWPW2*, the same test revealed that a model in which all species have the same *ω*is significantly less likely than one in which *ω* is elevated in lineages lacking a complete *PRDM9*; in particular, in canids and platypus (*p*=0.0003; **Table S10**). In fact, in canid lineages (fox and dogs), for which the loss of *PRDM9* is ancestral, *ZCWPW2* is no longer under any discernible selective constraint (testing a null model of *ω*=1, *p*=0.307; **Table S10**). Considered together with the observation that *ZCWPW2* is absent from all the other lineages in which a complete PRDM9 gene is clearly absent, these evolutionary analyses suggest that *ZCWPW2* is dispensable in the absence of a complete *PRDM9* ortholog.

The molecular function of ZCWPW2 is to our knowledge unknown. Like its paralog, it could be involved in the processing or repair of DSBs. If so, the observation that *Zcwpw1*-/- mice show defective DSB processing and repair [25-27] suggests that the role of ZCWPW2 cannot be completely redundant with that of its paralog. Alternatively, by reading the dual marks laid down by PRDM9, ZCWPW2 might help to recruit the recombination machinery (in particular SPO11) and thus play an earlier role in the positioning of DSBs. While in yeast, the link between histone modifications (specifically, H3K4me3) and the recruitment of Spo11 is made by Spp1 [46], in mammals, the ortholog of Spp1, CXXC1, is not essential for meiosis [47], and the gene that plays the analogous role has not yet been identified. Our analysis highlights ZCWPW2 as a potential candidate for this role, to be tested experimentally.

For *TEX15*, *ω* is also higher in lineages where *PRDM9* is absent or incomplete (*p*=0.0036 for fish and *p*=0.015 for mammals), but remains significantly below 1 (**Table S10**), indicating that in the absence of *PRDM9*, *TEX15* is not dispensable. If *TEX15* and *PRDM9* are indeed co-evolving, the relationship is likely to be indirect; for instance, recent work implicates TEX15 as an effector of piRNA-mediated transposable element (TE) methylation and silencing [48,49]. Male mouse knockouts of *Tex15* exhibit a meiotic arrest phenotype associated with the failure to repair DSBs and to undergo chromosomal synapsis [45], as well as the transcriptional activation of TEs [48,49]. This phenotype is similar to those observed in mouse knockouts of other piRNA-pathway genes, such as *Miwi* or *Dnmt3* [50]. In *Dnmt3* knockout mice, it has been shown that TEs accumulate both H3K4me3 marks and SPO11-dependent DSBs, suggesting that the methylation of TEs serves not only to silence them, but may also result in preventing their use as sites of recombination [50]. Thus, *TEX15* could conceivably play an important but indirect role in preventing the binding of PRDM9 to TEs.

For *FBXO47*, *ω* is higher in fish lineages where *PRDM9* is absent or incomplete (*p*=0.0023), but remains significantly below 1, while in mammals, there is no evidence for relaxation of constraint (**Table S10**). Like *TEX15*, if *FBXO47* and *PRDM9* are co-evolving, the relationship is likely to be indirect. *FBXO47* is a member of the F-box protein family, which act as recognition subunits of Skp1-Cullin1-F-Box protein (SCF) E3 ubiquitin ligase complexes [38,51]. Recently, *FBXO47* has been implicated as a key regulator of the telomere shelterin complex during meiotic prophase I, and in mice is necessary for telomere nuclear envelope attachment and subsequent events, including DSB repair [38]. One possibility for increased conservation of *FBXO47* in the presence of *PRDM9* would be if this role of FBXO47 contributes to the formation of a chromatin environment that aids in the repair of PRDM9-dependent DSBs, or possibly in the recruitment of ZCWPW1.

### Which loss came first?

While PRDM9 has two distinct roles––in specifying the location of DSBs and in facilitating their repair––the four candidate genes that we have identified may only be involved in one of these two roles. If so, the dependencies between the presence of PRDM9 and of these genes may be asymmetric. For instance, if we ignore possible pleiotropic roles of the candidate genes, and assume ZCWPW1 and FBX047 play roles in, or related to, repair but not DSB localization, we would predict that their loss occurs after that of *PRDM9* (as appears to have been the case in *Tachysurus fulvidraco* for both genes, and for *FBXO47* in *Xenopus laevis*; **Table S8B**). In contrast, if ZCWPW2 is involved in DSB localization but not repair, we would predict it could be lost before *PRDM9* (as was seemingly the case in two lineages of ray-finned fish; **Table S8B**). The phylogenetic data considered here do not allow us to distinguish between these scenarios: there is statistical evidence for a dependence of state transitions of *ZCWPW1*, *ZCWPW2, FBX047* and *TEX15* on *PRDM9* as well as vice versa (in all tests, maximum p<0.07, testing the null model of no dependence against either dependence as an alternative model; **Table S11**). These scenarios could potentially be distinguished by collecting more fine-grained phylogenetic information to pinpoint the specific lineages in which the first loss occurred, as well as in light of further experimental data.

### Outlook

Our phylogenetic analysis allowed us to identify novel putative interactors of PRDM9 that are promising candidates for functional studies. For this analysis, the power comes from the repeated losses of *PRDM9*––in our case, from eleven transitions from presence to absence. Confounding these kinds of analyses, however, are issues of data quality and in particular absences of complete *PRDM9* orthologs that reflect poor genome quality rather than true losses. To address this issue, we validated any absence in Refseq with whole genome searches and where possible, *de novo* assemblies from RNA-seq data, leading us to realize that in one case (*MEI1*), the apparent co-evolution with *PRDM9* was in fact spurious.

A more subtle but related issue stems from a phylogenetic signal of genome quality, which can lead to apparent clustering of losses. To minimize this issue, we restricted our analysis to genomes that included most “core” eukaryotic genes (**Fig. S7**) and downsampled our tree to include at most three species below every inferred *PRDM9* loss. As genome qualities improve and as their assemblies become more uniform (eg., [52]), these issues should be alleviated. Moreover, as species are added to the phylogeny, additional losses will be identified: as one example, our identification of two species of frogs with a complete *PRDM9* revealed that *PRDM9* had not been lost once in the common ancestor, as had been inferred using fewer species by Baker et al. (2017), but has instead been lost more than once within amphibians. This discovery also suggests that frogs may be an interesting clade within which to study the steps by which PRDM9 and its partners are lost.

Beyond the application to *PRDM9* and meiotic recombination, our analysis illustrates how long-standing phylogenetic approaches can now be applied to comparative genomic data to identify novel molecular interactions [53]. Such analyses need not be restricted to measurements of presence or absence of whole genes, as we have done here, but could focus exclusively on specific domains, indicative of specific subfunctions, or consider how rates of evolution in specific domains depend on the presence or absence of other genes. With the explosion of high quality and more representative sets of genomes now coming on line (e.g., [52], [54]), and the development of statistical methods that consider both binary and continuous character evolution jointly, we expect this type of approach to become increasingly widespread.

## Material and Methods

### 1. Identification of PRDM9 orthologs

As a first step towards characterizing the distribution of *PRDM9* in vertebrates, we identified putative *PRDM9* orthologs in the RefSeq database with a *blastp* search [30], using the N-terminal portion of the *Homo sapiens* PRDM9 protein sequence containing KRAB, SSXRD and SET domains as the query sequence (RefSeq accession: NP_001297143; amino acid residues 1-364). We downloaded the corresponding GenBank file for 5,000 hits (3,400 unique genes from 412 species) and characterized the presence or absence of KRAB, SSXRD and SET domains for each record using the Conserved Domain, Protein Families, NCBI curated and SMART databases (CDD [55]; Pfam (REF); NCBI curated (REF); SMART (REF); accessions cl02581 and cl09744 for the KRAB and SSXRD domains respectively, and accessions cl40432 and cl02566 for the SET domain), annotating each domain as present if that domain had an e-value less than 1 in any of the four databases. We then removed alternative transcripts from the dataset by preferentially keeping, for each unique gene, the transcript with the maximal number of annotated domains. When there were multiple transcripts with the same maximal number of domains, we kept the longest one.

Because *PRDM9* shares its SET domain with other PRDM family genes and its N-terminal domains with members of the KRAB-ZF and SSX gene families, many of these hits are potential PRDM9 paralogs. To identify bona fide *PRDM9* orthologs from this initial set of genes, we sought to build phylogenetic trees specific to the KRAB, SSXRD, and SET domains and remove homologs that cluster with genes annotated as distantly related paralogs of *PRDM9*. To this end, we extracted the amino acid sequences for complete KRAB, SSXRD, and SET domains, and for each domain, constructed neighbor-joining trees using Clustal Omega [56]. Utilizing the KRAB and SSXRD domain-based trees, we identified and removed 87 genes that visually cluster with members of the SSX gene family (**Fig. S1A-B**). Analyzing the SET domain-based tree, we identified and removed 2,637 genes that group with other members of the *PRDM* gene family (**Fig. S1C**; see figure legend for details). We ultimately retained 625 genes, each of which cluster with *PRDM9* in one or more of these trees.

By this approach, in the 412 species considered, we identified 209 *PRDM9* orthologs containing KRAB, SSXRD and SET domains from 155 species, as well as 13 *PRDM9* orthologs containing KRAB and SET domains for which we were unable to detect an SSXRD domain with an e-value less than 1 from an additional 11 species. For the 246 species for which we were unable to identify a *PRDM9* ortholog spanning KRAB and SET domains in our initial search of the RefSeq database, we sought to verify that *PRDM9* was truly absent using a number of approaches.

As a first step, we performed an additional blastp search against the non-redundant protein sequence (nr) database, targeting only those species in order to identify any annotated gene record missed in our initial search of the RefSeq database. We downloaded the corresponding GenBank file for each hit with >55% coverage and >40% identity and, after removing records corresponding to those we had previously identified, annotated domains and removed alternative transcripts as before. We then verified the orthology of the remaining records by blasting each protein sequence against the human RefSeq database, accepting it as a PRDM9 ortholog if the top hit was PRDM9 or its paralog PRDM7. This approach enabled the identification of an additional 9 PRDM9 orthologs, including one containing KRAB, SSXRD and SET domains, and one containing KRAB and SET domains.

Next, we performed a series of *tblastn* searches of the whole genome of the 244 species remaining using the N-terminal portion of the *Homo sapiens* PRDM9 protein as a query. When we were unable to retrieve any promising hits with the human protein sequence, we re-performed the *tblastn* search using the N-terminal portion of a *PRDM9* ortholog from a species closely related to the focal species. In order to identify which of the identified contigs corresponded to genuine *PRDM9* orthologs (as opposed to paralogs such as *PRDM11*), we performed blastp searches against the *Homo sapiens* RefSeq database using the aligned protein sequences as query sequences. Contigs containing the relevant alignments spanning KRAB and/or SET domains were then downloaded and the aligned region including 10,000 of flanking sequence was extracted and input into *Genewise* [57], using the PRDM9 protein sequence from *Homo sapiens* or a closely related species as a guide sequence (see **Table S2** for details). In genomes from 10 species, we identified separate contigs containing the KRAB domain and the SET domain. In these cases, the contigs were concatenated before use as input in *Genewise*. These approaches enabled us to identify an additional 53 *PRDM9* orthologs from 33 species, including 21 *PRDM9* orthologs containing KRAB, SSXRD and SET domains from 21 species, and 24 *PRDM9* orthologs containing KRAB and SET domains but for which we were unable to identify the SSXRD domain from 11 species.

These analyses left 210 species for which we were unable to identify a *PRDM9* ortholog with both KRAB and SET domains. For these species, with the exception of 94 birds and crocodiles and 78 percomorpha fish, we additionally searched testis RNA-seq datasets when possible, including those generated for this study (see below; **Table S3**); this approach enabled us to identify two additional *PRDM9* orthologs containing KRAB and SET domains from two species of fish.

From this analysis, and given the phylogenetic relationships among species given by the TimeTree tool [35], we inferred 20 putative complete or partial losses of PRDM9 across the 412 species represented in the RefSeq database. Of these, 7 losses were supported by the absence of PRDM9 in two or more closely related species: in percomorpha and beryciformes fish, characiformes and siluriformes fish, cypriniformes fish, polypteridae fish, frogs, birds and crocodiles, and canids. The remaining 13 inferred losses each corresponded to an individual species. In order to identify whether or not any of these 13 latter absences could be supported by additional species, and to more accurately infer the dates of each loss, we sought to investigate the status of PRDM9 in species closely related to each putative loss event. To this end, we investigated the whole genomes of an additional 18 species and RNA-seq datasets from an additional 4 species as before, with one species represented by both a whole genome sequence and a corresponding RNA-seq dataset (*Ambystoma mexicanum*). This approach enabled us to identify an additional 6 PRDM9 orthologs containing KRAB, SSXRD and SET domains from 6 species, as well 15 additional species putatively lacking a complete PRDM9 gene. In doing so, we found that 2 additional losses were supported by the absence of PRDM9 in two or more closely related species: in osteoglossomorpha fish, as well as a loss within lizards shared by *Anolis carolinensis* and *Scleropages formosus*. For each of the 11 remaining instances in which only a single species was found to be lacking PRDM9, the most closely related species considered possessed a complete PRDM9 ortholog. While independent work supports our finding of a partial loss of PRDM9 in platypus (J. Hussin and P. Donnelly, personal communication), we do not have confirmatory evidence of absence for the remaining 10 species, and therefor treat these species as having an uncertain PRDM9 status. Moreover, we identified two species of frogs carrying complete PRDM9 orthologs. This discovery suggests that PRDM9 has been lost repeatedly within amphibians – at least once in salamanders, and at least three times within frogs (with each of these four putative loss events being supported by the absence of the PRDM9 in two or more closely related species).

Lastly, we include in the list of species considered an additional 13 species for which we had previously identified complete PRDM9 orthologs [11] but which were not directly examined here (**Table S4**). Altogether, this pipeline resulted in the identification of 193 species in which we find a complete *PRDM9* ortholog containing KRAB, SSXRD and SET domains, 26 species for which we identify *PRDM9* orthologs containing KRAB and SET domains but not SSXRD domains, 218 species for which we have evidence for the absence of a complete *PRDM9* gene, and 9 species for which we were unable to make a confident determination (see **Tables S1-S4,** **Figure 1**).

For each of the *PRDM9* orthologs that we identified, we characterized the conservation of three key tyrosine residues that have been shown to underlie the catalytic function of the human SET domain in vitro (i.e., Y276, Y341, and Y357; [58]) and for Y357, in vivo in mouse [10]. To this end, we constructed an alignment of the SET domain using Clustal Omega [56] and extracted the residues aligning to the human tyrosine residues from each of 678 SET domains (**Table S1**).

### 2. Verification of genomic calls using RNA-seq data

For four species in which we identified no *PRDM9* ortholog or only a partial ortholog, we investigated whether a complete *PRDM9* ortholog may nonetheless be present using RNA-seq data. We therefore sought to verify its absence from *Anolis carolinensis,* a species in which we had been unable to find a *PRDM9* ortholog in the genome assembly or Refseq, as well as a second reptile species, *Sceloporus undulatus,* for which Refseq data and a genome sequence were not available. To this end, we built a *de novo* RNA transcriptome assembly and tested for the expression of PRDM9 in testis and other tissue samples (see below).

Similarly, in two species in which we had originally identified only a partial ortholog of *PRDM9* (*Astyanax mexicanus* and *Clupea harengus*), we wanted to verify the incomplete domain structure inferred from the genome sequence by conducting a *de novo* transcriptome assembly (in *Clupea harengus*, this analysis turned out to be unnecessary, as an updated reference genome, GCA_000966335.1, contains a complete *PRDM9*). To this end, we analyzed RNA-seq data from male gonad surface from *Astyanax mexicanus* and liver and testis from *Clupea harengus*.

Dissected tissue samples preserved in RNAlater were kindly provided to us by Arild Folkvord and Leif Andersson (*Clupea harengus*), Cliff Tabin (*Astyanax mexicanus*), Tonia Schwartz and Tracy Langkilde (*Sceloporus undulatus*), and Athanasia Tzika (*Anolis carolinensis*). These samples were stored at -20°C until extraction and library preparation. Total RNA was extracted using the Qiagen RNeasy kit (Valencia, CA, USA) following the manufacturer’s protocol. RNA was quantified and assessed for quality on a Qubit fluorometer and approximately 1 μg of total RNA was input for library preparation using the Kapa RNA-seq kit. Samples were prepared following the manufacturer’s protocol, except that half reactions were used. Briefly, mRNA was purified using manufacturer’s beads and chemically fragmented. First and second-strand cDNA was synthesized and end-repaired. Following A-tailing, each sample was individually barcoded with an Illumina index and amplified for 12 cycles. In order to evaluate the library quality and size distribution, libraries were evaluated on an Agilent Tapestation. The libraries were then sequenced over two runs on the NextSeq 550 at Columbia University to collect paired-end 150 bp reads.

Illumina sequencing reads (248,820,547 2x150 base pair (bp) paired-end reads) were demultiplexed into individual sample fastq files with the software bcl2fastq2 (v2.20.0, Illumina). The FastQC software [59] was used for visual inspection of read quality. Adapters and low-quality reads were trimmed with the Trimmomatic software, which is bundled as a plugin within the Trinity *de novo* assembler [60] (v2.8.5) and was enabled using the *--trimmomatic* flag. The default trimming settings (phredscore>=5; slidingwindow:4:5; leading:5, trailing:5; minlen:25) were used following [61] recommendations. The pair-end reads were trimmed and *de novo* transcriptomes assembled with Trinity (v2.8.5) using the following parameters: --seqType fq --SS_lib_type FR -- max_memory 100G --min_kmer_cov 1 --trimmomatic --CPU 32. Details on assembly quality are shown in **Fig. S2**. Gene expression data for all four species (*Anolis carolinensis, Sceloporus undulatus, Clupea harengus, Sceloporus undulatus*) are available from the NCBI sequence read archive (Bioproject PRJNA605699, SRA accessions: SRR11050679-SRR11050687).

To evaluate whether PRDM9 was present in the transcriptome data, we conducted a *tblastn* search (e-value ≤ 1e-5) against each *de novo* assembly using the human PRDM9 protein sequence (without its rapidly evolving zinc finger array) as a query, and we classified the domain presence of up to five top hits using CDD blast [55]. For a given species, if the KRAB and SET domains were not identified in any transcript, *PRDM9* was considered incomplete. The inability to identify PRDM9 could indicate either that the gene is not expressed or that we lack the appropriate cell types or sequence coverage to detect it. To assess our power to detect PRDM9 from the testis RNA-seq data, we followed methods outlined in [11]. Specifically, for each transcriptome, we evaluated whether we could identify transcripts from six genes with highly conserved roles in meiotic recombination [62] (*HORMAD1*, *MEI4*, *MRE11A*, *RAD50*, *REC114*, and *SPO11*). To identify the transcripts orthologous to each of these genes, we performed a *tblastn* search (e-value ≤ 1e-5) of the *Homo sapiens* reference protein sequence against each *de novo* transcriptome. We considered PRDM9 to be absent if we detected expression of all six genes but not a complete *PRDM9*; by these criteria, we found PRDM9 to be missing from *A. carolinensis*, *S. undulatus*, and *A. mexicanus*.

Using the same approach to *de novo* assembly and gene detection, we also analyzed publicly available RNA-seq datasets from testis for 28 additional species (**Table S3**), either to verify the absence of PRDM9 (see above) or of one of the candidate genes (see below).

To estimate the expression levels of the Trinity-reconstructed transcripts, we used RSEM [63] (v1.3.1) implemented through Trinity (v2.8.5). We first aligned the RNA-seq reads from each sample to the newly generated *de novo* assembled transcriptome (see above) using the alignment method bowtie [64] (v1.2.2). We then extracted quantification information for each gene of interest from the RSEM output (in fragments per kilobase of transcript per million mapped reads or FPKM) (**Fig. S3**).

### 3. Choice of candidate genes and orthology assignments

To identify a set of genes that may co-evolve with *PRDM9*, we relied on three publicly available datasets, namely: (i) 39 genes associated with variation in recombination phenotypes in a genome-wide association study in humans [33]. Of the variants reported to be associated with recombination phenotypes, six were found in intergenic regions; we included the subset of two cases in which the authors assigned these variants to nearby genes (*ZNF84* and *ZNF140*). (ii) 193 genes co-expressed with PRDM9 in single cell data from mouse testes. Specifically, we considered the top 1% of genes based on their gene expression loadings in component 5, the component in which PRDM9 has the highest loading [31]. (iii) 36 genes known to have a role in mammalian meiotic recombination based on functional studies [32].

Genes co-expressed with PRDM9 in mouse spermatogenesis were converted to human gene symbols using the package biomaRt in R [65]. Fifteen of these genes did not have an orthologous human gene symbol (*Gm7972*, *H2-K1*, *Gm4349*, *Ddx43*, *Atad2*, *Xlr4c*, *Gm364*, *Tex16*, *4933427D06Rik*, *AI481877*, *H2-D1*, *Trap1a*, *Xlr4a*, *2310035C23Rik*, and *Tmem5*) and eight other genes mapped to more than one human gene symbol (*Msh5*, *Cbwd1*, *Nxf2, Cbwd1*, *Fam90a1b*, *Srgap2*, *Cdk11b*, *Gm15262*). Keeping all mapped gene symbols yielded 185 genes; combined with the two other sources, 241 genes were tested for their co-evolution with *PRDM9* (**Figure S6**). A supplementary file describing each meiosis candidate gene is available in **Table S5**.

For the 241 genes, we characterized whether the ortholog is present in its complete form across vertebrate species. To this end, we first downloaded all the vertebrate RefSeq protein sequences available on the NCBI database (accessed on June 3, 2020), corresponding to 339 species. Of these, we filtered out 32 species that were missing 10 or more BUSCO core genes (out of a total of 255 genes) [66], reasoning that their genomes were sufficiently incomplete that they may be missing orthologs by chance (see **Figure S7**). Of the remaining 307 species, we further excluded 29 species in order to remove polytomies observed in the phylogeny; specifically, we removed the minimal number of species necessary to remove each polytomy while preserving any transitions in the state of *PRDM9*. Moreover, to minimize possible phylogenetic signals generated by genome assembly quality, we thinned the tree such that for each *PRDM9* loss along the phylogeny, we kept at most three species representing that loss. In cases where a loss was ancestral to more than three species in our dataset, we picked three distantly related species with the best genome assemblies, as measured by the BUSCO score. In the end, we retained 189 species: 134 mammals, 3 birds, 6 amphibians, 18 reptiles, 2 percomorph fish, 3 cypriniformes fish, 20 other ray-finned fish, 2 cartilaginous fish, and one jawless fish. This phylogeny includes representative species for 11 of the 13 inferred PRDM9 losses: species of *Bufonidae* frogs and salamanders were not included due to the absence of available gene annotations; also due to the lack of gene annotations for frog species with PRDM9, within these 189 species, the losses in *Xenopus* and *Dicroglossidae* frogs cannot be distinguished from a single event.

For each candidate gene in each species, we performed a blastp search of the human ortholog against the RefSeq database of the species and kept up to five top hits obtained at an e-value threshold of 1e-5. We inferred the domain structure of each hit using the Batch CD-Search [67], and considered a domain as present in a species if the e-value was ≤ 0.1. We considered genes to be complete orthologs if they contained the superfamily domains found in four representative species of the vertebrates phylogeny carrying a complete PRDM9 (*Esox lucius* (fish) *Geotrypetes seraphini* (caecilian), and *Pseudonaja textilis* (snake)), at an e-value threshold of 1e-4. For the 15 genes (*FANCB*, *FMR1NB*, *GPR137C*, *HAUS8*, *M1AP*, *MEI1*, *SPATA22*, *CLSPN*, *FBXO47*, *HMGA2*, *HSF2BP*, *IQCB1*, *LRRC42*, *PRAME*, *SYCE2*) in which no detectable domains were present, we annotated the presence or absence of the gene using the blastp results alone. In the end, we built a matrix of presence or absence across species and candidate genes to be used in the phylogenetic test (see **Table S6**).

### 4. Testing for the co-evolution of PRDM9 and candidate genes

To test for the co-evolution of *PRDM9* and each candidate gene, we need to account for the phylogenetic relationships among the species considered. To obtain these relationships and time-calibrated branch lengths, we used the TimeTree resource (http://timetree.org/; [35], accessed on June 10, 2020). Of the 189 species included in the phylogenetic tests, 9 were not present in the TimeTree database; in those cases, we used information from a close evolutionary relative to determine their placement and branch lengths.

For this test, we consider *PRDM9* as present if it contains KRAB and SET domains or incomplete/missing if one of those domains is absent (**Tables S4** and **S8**). We do not rely on the SSXRD domain when making these calls because its short length makes its detection at a given e-value threshold unreliable. Notably, for 19 of the 26 species with *PRDM9* orthologs containing KRAB and SET domains, but not SSXRD domains with an e-value < 1, we are able to detect the SSXRD domain when using an e-value threshold of 1000 (**Table S1**). We additionally do not rely on the ZF array because its repetitive nature makes it difficult to sequence reliably.

We tested whether state changes of intact candidate genes were unexpectedly coincident with state changes of the intact *PRDM9* using the software *BayesTraitsV3* [68]. We did so by comparing the statistical support for two models: a null model in which *PRDM9* and a given candidate gene evolve independently of one another along the phylogeny versus an alternative model in which the gain (“1”) and loss (“0”) of a gene is dependent on the status of *PRDM9* and vice versa. We compared the likelihoods of the two models using a likelihood ratio test with 4 degrees of freedom, and reported a p-value uncorrected for multiple tests (**Table S7**). For each gene and model, 100 maximum likelihood tries were computed and the maximum likelihood value was retained. A quantile-quantile plot was drawn to access the distribution of p-value, and the R package “Haplin” was used to compute pointwise confidence intervals (CI). To control for the false discovery rate (FDR), we computed q-values using the R package “qvalue” and set a 50% FDR threshold.

Given the phylogenetic distribution of *PRDM9*, it is likely that a *PRDM9* ortholog was present in the common ancestor of vertebrates [11,12]. Based on this prior knowledge, we restricted the state of *PRDM9* at the root of the phylogeny to always be present. In turn, for each candidate gene, we set a prior in which it had 50% probability of being present and 50% probability of being absent. We also used this prior for the state of *PRDM9* in the 9 species that lack *PRDM9* but where the loss was not supported by a closely related species (i.e., for which we considered the status uncertain).

For *FBXO47, TEX15*, *ZCWPW1* and *ZCWPW2*, we also explored restrictions on the rates in the dependent model, such that their state transitions depend on PRDM9 (model X) or the state transitions of *PRDM9* depends on theirs (model Y), rather than both being true. For these tests, we compared the likelihoods of each dependent model against our independent null model using a likelihood ratio test with 2 degrees of freedom. For each gene and model, 100 maximum likelihood tries were computed and the maximum likelihood value was retained.

We also explored whether redefining a complete PRDM9 ortholog as containing not only the KRAB and SET domain but also the SSXRD domain would change the statistical significance. By using the improved calls (see below), only *ZCWPW2* remains significant (*p*=0.004) and *ZCWPW1* marginally so (*p*=0.056) (**Table S9B-C**).

### 5. Improving gene status calls of top candidate genes

For the five genes with a FDR ≤ 50% (**Figure 2A**), we sought to improve our calls by building phylogenetic trees based on domains in the genes and examining the clustering patterns visually, as well as by searching for orthologs in whole genome assemblies and testis transcriptomes (following the same procedures described for *PRDM9*). These improved calls were then used to rerun the phylogenetic independent contrast tests, following the same implementation as previously; the p-values for these improved gene models are shown alongside the original ones in **Table 1**. Below we provide an overview of the steps we took for each candidate gene. For each gene we provide descriptions of identified orthologs and how they were identified in **Table S1**, specific details about orthologs identified from whole genome assemblies in **Table S2**, our improved calls per species in **Table S8A**, and a summary of loss events in **Table S8B**.

#### i. MEI1

For *MEI1*, an initial blastp search of the vertebrate RefSeq database using the human sequence as query resulted in the identification of 422 *MEI* orthologs from 372 species. We note that, for *MEI1*, we did not find any domain annotations, and therefore did not perform phylogenetic analysis to support the identification of these orthologs. However, each homolog identified in our initial RefSeq analysis was annotated as either *MEI1* or *MEI1-like*. We thus labeled each species as having a complete ortholog if an ortholog was present. This approach resulted in the identification of *MEI1* ortholog for 187 of the 189 species used for our co-evolutionary test. For the remaining 2 species, we sought to identify *MEI1* orthologs from whole genome sequences following the same procedures described for *PRDM9*. This approach allowed us to identify a *MEI1* ortholog in every species, revealing that in fact, *MEI1* has not been lost among the vertebrate species examined (**Tables S1** and **S2**).

#### ii. ZCWPW1 and ZCWPW2

Because *ZCWPW1* and *ZCWPW2* are paralogs, we performed our analyses of these genes together. To this end, we combined the datasets of genes identified in our initial RefSeq blastp search to create a dataset of 977 putative orthologs from 363 species. We then extracted amino acid sequences and built neighbor-joining trees using Clustal Omega for both the zf-CW and PWWP domains ([56]; accessions cl06504 and cl02554; **Figure S9A-B**). Utilizing these trees, we removed 573 genes that visually clustered with genes annotated as distantly related paralogs, such as members of the MORC and NSD gene families. We additionally relied on these trees to more confidently label which genes were *ZCWPW1* orthologs and which were *ZCWPW2* orthologs based on where they clustered in the tree. We considered orthologs as complete if they contain both PWWP and zf-CW domains with e-values < 1. This approach resulted in the identification of 193 complete *ZCWPW1* orthologs from 188 species, and 187 complete *ZCWPW2* orthologs from 180 species.

Among the 189 species used in our co-evolutionary test, 164 had complete *ZCWPW1* orthologs and 154 had complete *ZCWPW2* orthologs on the basis of this initial search. For the 25 species missing a complete *ZCWPW1* ortholog, and for the 35 missing a complete *ZCWPW2* ortholog, we sought to identify the orthologs from whole genome sequences following the same procedures as described for *PRDM9*. This approach enabled us to identify an additional 3 complete *ZCWPW1* orthologs from 3 species, and an additional 11 complete *ZCWPW2* orthologs from 11 species (**Tables S1** and **S2**). For the remaining species, we checked the putative loss of *ZCWPW1* or *ZCWPW2* using RNA-seq data when available, which led to the identification of an additional 2 complete *ZCWPW1* orthologs, but no additional *ZCWPW2* orthologs (**Tables S1** and **S3**). We additionally added one ZCWPW1 ortholog from the common shrew (*Sorex araneus*) from the Ensemble database, which had been identified previously but was absent from NCBI [26,27]. Lastly, we sought to identify *ZCWPW2* from the whole genome sequence of a species of frog with PRDM9 (*Ranitomeya imitator*) not otherwise included in our co-evolutionary test in order to distinguish whether or not the absence of *ZCWPW2* in *Xenopus* and *Dicroglossidae* frogs corresponded to a single loss or multiple events. We were able to identify *ZCWPW2* from this species, suggesting that *ZCWPW2* has been lost multiple times within frogs (**Tables S1**, **S2** and **S8**).

#### iii. TEX15

For *TEX15*, our initial blastp search resulted in the identification of 900 putative orthologs from 363 species. We similarly utilized a tree built using the DUF3715 domain to remove 667 genes that cluster with distantly related paralogs, in particular, *TASOR* and *TASOR2* (**Figure S9C**). When making our final calls about *TEX15* orthologs, we labeled them as complete if they contained both DUF3715 and TEX15 domains (accessions pfam12509 and pfam15326). This approach resulted in the identification of 179 complete *TEX15* orthologs from 175 species. Among the 189 species used for our co-evolutionary test, 150 had complete *TEX15* orthologs on the basis of this initial search. For the 39 species missing a complete *TEX15* ortholog, we sought to identify the orthologs from whole genome sequences, following the same procedures as described for *PRDM9*. In this way, we identified an additional 29 complete *TEX15* orthologs from 28 species (**Table S2**). For the remaining species, we checked if we could find *TEX15* using RNA-seq data, when available, and found one additional complete *TEX15* ortholog by this approach (**Tables S1** and **S3**).

#### iv. FBXO47

For *FBXO47*, an initial blastp search of the vertebrate RefSeq database using the human sequence as query resulted in the identification of 386 putative *FBXO47* orthologs from 380 species. We did not perform phylogenetic analysis to support the identification of these orthologs: While we detected a domain (F-BOX) in the human *FBXO47* gene, due to its high e-value in humans (e-value = 0.01), we did not rely on its presence or absence when inferring the whether or not a complete *FBXO47* gene was present in each species. However, each homolog identified in our initial RefSeq analysis was annotated as either *FBXO47* or *FBXO47*-like with the exception of one *CWC25* gene, which was removed. We thus labeled each species as having a complete ortholog if an ortholog was present. This approach resulted in the identification of *FBXO47* ortholog for 181 of the 189 species used for our co-evolutionary test. For the remaining 8 species, we sought to identify *FBXO47* orthologs from whole genome sequences and/or RNA-seq datasets following the same procedures described for *PRDM9*; however, we were unable to identify any additional FBXO47 in this way (**Tables S1** and **S3**).

### 6. Conservation of residues in *ZCWPW2*

We carried out a residue conservation analysis using an approach proposed by [69], using code *score_conservation.py* available at https://compbio.cs.princeton.edu/conservation/. This approach quantifies the Jensen–Shannon divergence between the amino acid distribution of the focal residue and a “background amino acid distribution.” The alignment of *ZCWPW2* was produced using Clustal Omega (using default parameters) within MEGA (version 7, [35,70]). As recommended, the overall background amino acid distribution was drawn based on the BLOSUM62 amino acid substitution matrix provided by the software [69]. Any column of the gene sequence alignment with more than 30% gaps was ignored. A window size of 3 was used to incorporate information from sequential amino acids, as recommended by the default settings.

### 7. Evidence for relaxed selective constraint in the absence of *PRDM9*

To test for possible relaxed selection in species without a complete *PRDM9*, we used the program *codeml* within PAML [71,72]. *Codeml* uses protein coding sequences to estimate the ratio of non-synonymous to synonymous substitution rates (*ω* = *d_N_*/*d_S_*). Values of *ω*significantly less than 1 are indicative of purifying selection, i.e., of the functional importance of the gene.

To this end, we considered each major clade (fish, mammals, reptiles, amphibians) separately and extracted and aligned coding nucleotide sequences from NCBI for multiple species. We aligned those sequences in a codon-aware manner using Clustal Omega (using default parameters) within MEGA (version 7, [35,70]) and inspected the codon-aware alignment visually to ensure that the same isoforms were used across species. For each multi-species alignment, we tried two approaches: (i) We estimated *ω* under a null model assuming the same *ω* across all branches and an alternative model in which there are two *ω* allowed: one *ω* value in species with a complete *PRDM9* and a second *ω* for the branches in which *PRDM9* is absent or incomplete (including the internal branches on which *PRDM9* may have been lost); (ii) We considered the same null model with the same *ω* across all branches; and an alternative model with one *ω* value in species with a complete *PRDM9*, a second *ω* for the branches in which *PRDM9* is absent or incomplete and additional *ω* values for each branch on which *PRDM9* was inferred to be lost (a different one for each independent loss, as the *ω*value averaged over the branch will depend on when along the branch *PRDM9* was lost). For (i), significance was assessed using a likelihood ratio test with 1 degree of freedom; for (ii), by the number of degrees of freedom corresponded to the number of distinct *ω* values minus 1. If *ω* values were found to be significantly higher in species without a complete *PRDM9*, we tested whether or not we could reject *ω* = 1 for these species. For two cases in which we could not obtain a multi-species alignment that included the whole coding sequence (*ZCWPW1* in fish and *TEX15* in amphibians), we instead used the pairwise model (runmode: -2 within PAML) on alignments for a pair of species, and tested whether we could reject *ω* = 1 for species lacking *PRDM9* by comparing a model allowing *ω* to vary versus a null model fixing the *ω* value at 1, with 1 degree of freedom.

## Acknowledgments

The authors would like to thank Arild Folkvord and Leif Andersson (*Clupea harengus*), Cliff Tabin (*Astyanax mexicanus)*, Tonia Schwartz, and Tracy Langkilde (*Sceloporus undulatus*), and Athanasia Tzika (*Anolis carolinensis*) for providing testis, muscles, and liver samples, and Mackenzie Keegan for extraction and library preparation of samples. The authors are also grateful to Andres Bendesky for use of his lab space and sequencing facilities and to Andrew Meade for helpful discussions about the software *Bayestraits*. In addition, we thank Todd Jackman and Daniel Wells for pointing out mistakes in the previous version of our manuscript, now corrected.

## Data and Code availability

The following data sets were generated

1. **RNA-seq data for two fish species and two reptile species (fastq files)** Cavassim M I A, Baker Z, Hoge C, Schierup M, Schumer M and Przeworski M (2020) Available from the NCBI BioProject (accession number: PRJNA605699).
2. **Code availability** Code generated for this study can be found at https://github.com/izabelcavassim/PRDM9_analyses
3. **Supplementary material** Supplementary material is available on https://www.dropbox.com/sh/pihq6a643fz21js/AAANGJWpALT42MCrrsJdlSUHa?dl=0

## Author contributions

**Conceptualization:** M.I.A.C., Z.B., and M.P.; **Methodology:** M.I.A.C., Z.B., C.H., M.S., and M.P.; **Formal Analysis:** M.I.A.C., Z.B.; **Investigation:** M.I.A.C., Z.B., C.H., M.S.; **Resources:** M.H.S., M.S., M.P.; **Data curation:** M.I.A.C., Z.B., M.S., and M.P.; **Writing - Original Draft:** M.I.A.C. Z.B., M.S., and M.P. **Writing - Review and Editing:** M.I.A.C., Z.B., C.H., M.S., and M.P; **Visualization:** M.I.A.C. and Z.B. **Supervision:** M.H.S., M.S., and M.P. **Project administration:** M.P.; **Funding acquisition:** M.H.S., M.S., and M.P.

## Supplementary tables

**Supplementary Table 1.** Genes identified in this study. For *PRDM9* and each candidate gene that was initially identified as significantly coevolving with *PRDM9* (*ZCWPW1, MEI1, ZCWPW2, TEX15*, and *FBX047*), we provide a table detailing, for each ortholog that we identified, which species it is from, how we identified it, its inferred domain architecture, its amino acid sequence, as well as various details about these domains (including their coordinates, e-values, and sequences). For *PRDM9* orthologs, we additionally report which amino acid residues align to three catalytic tyrosine residues in the human SET domain.

**Supplementary Table 2.** Description of genes found from whole genome sequences. For *PRDM9* and each gene initially identified as significantly coevolving with *PRDM9* (*ZCWPW1, MEI1, ZCWPW2*, and *TEX15*), we provide a table detailing where and how each ortholog identified in our analysis of whole genome assemblies was obtained. *FBXO47* is excluded from this table because no *FBXO47* orthologs were identified from whole genome sequences.

**Supplementary Table 3.** Description of species for which a de novo assembly of testis transcriptomes was generated in order to verify the structure and expression of *PRDM9* and four significant genes (ZCWPW1, ZCWPW2, TEX15 and FBX047). To this end, we used publicly available RNA-seq data (downloaded from NCBI) [75] and in a subset of cases indicated with a star, generated our own data which are available from the NCBI sequence read archive (Bioproject PRJNA605699, SRA accessions: SRR11050679-SRR11050687, see Methods for further details).

**Supplementary Table 4.** The distribution of *PRDM9* orthologs across 446 vertebrate species. For each species, we describe how many *PRDM9* orthologs we identified of each unique domain architecture, the domain architecture of the most complete *PRDM9* ortholog from that species, and whether any *PRDM9* ortholog from that species with the most complete domain architecture has conserved three catalytic tyrosine residues in the SET domain. We additionally include columns comparing these results to those previously described in Baker et al. 2017, noting instances where we have revised our calls of domain architecture.

**Supplementary Table 5.** Description of candidate genes used in the phylogenetic tests. The 241 genes selected for the tests were based on three different sources: (1) genes most highly co-expressed with *PRDM9* in mouse testis single cell analyses [31], (2) genes associated with variation in recombination phenotypes in humans [33], and (3) genes known to have a role in mammalian meiotic recombination from functional studies (as summarized in the review by [32]). The genomic coordinates (column named ‘Start position’) of each gene were based on the GRCh38/hg38 human reference. The function of each candidate gene is described based on the definition from its source.

**Supplementary Table 6.** Presence and absence matrix computed for all candidate genes used in phylogenetic tests for coevolution with *PRDM9* (139 genes). We defined a gene as complete (“1”) when it contained all the domains observed in four representative vertebrate species with a complete *PRDM9* sequence, and incomplete (“0”) if the gene was not detected in the Refseq database or if it did not include all the domains shared across four species (see Methods for details).

**Supplementary Table 7.** Phylogenetic tests and p-values. P-values were computed by evaluating the patterns of presence or absence of *PRDM9* across 189 vertebrates against the patterns of presence or absence of candidate genes. Two models were tested using BayestraitsV3 [36]: a null model in which *PRDM9* and a given candidate gene evolve independently of one another along the phylogeny versus an alternative model in which the gain (“1”) and loss (“0”) of the candidate gene is dependent on the status of *PRDM9* and vice versa. See Pagel, 1994 and the BayesTraitsV3.0.2 manual for further discussion of these models and rates description.

**Supplementary Table 8.** The distribution of *PRDM9* and four genes initially found to be significantly coevolving with *PRDM9* (*ZCWPW1, ZCWPW2, TEX15*, and *FBXO47*) across 189 vertebrate species. MEI1 is not considered because in the curation of the calls, it was found to be present in all species (see text). (A) Curated calls for the presence or absence of complete genes based on searches of RefSeq, whole genome assemblies, and RNA-seq data (see **Tables S1- S3**). We additionally include the most complete domain architecture of orthologs from each species for each gene. (B) Summary of losses inferred for *PRDM9*, *ZCWPW1, ZCWPW2, TEX15* and *FBXO47* among the 189 vertebrate species used in our co-evolutionary test.

**Supplementary Table 9.** (A) Results of phylogenetic tests when considering the pairwise co-evolution of the candidate genes with each other. (B) Results of phylogenetic tests when considering the SSXRD domain in *PRDM9* classification. P-values were computed by evaluating the patterns of presence or absence of *PRDM9* across 189 vertebrates against the patterns of presence or absence of candidate genes. Two models were tested using BayestraitsV3 [36]: a null model in which *PRDM9* and a given candidate gene evolve independently of one another along the phylogeny versus an alternative model in which the gain (“1”) and loss (“0”) of a gene is dependent on the status of *PRDM9* and vice versa. See [37] and the BayesTraitsV3.0.2 manual for further discussion of these models and rates description. (C) Results of phylogenetic tests when considering the SSXRD domain in *PRDM9* classification and the curated calls for *ZCWPW1, ZCWPW2, TEX15*, and *FBXO47*.

**Supplementary Table 10.** Tests for differences in the rates of amino acid evolution in three significant genes (*ZCWPW1, ZCWPW2, TEX15* and *FBX047*) between representative species with and without a *PRDM9* ortholog. To determine whether species lacking a *PRDM9* ortholog showed evidence for relaxed selection pressures in co-evolving genes, we estimated *ω*(dN/dS) using the Branch model within PAML [76] under two models: a null model assuming the same *ω* across all branches of the phylogeny, and an alternative model in which there are two *ω* values allowed: one *ω* value in species lacking a functional *PRDM9* and a second *ω* for the rest of the branches. The clades evaluated in each test are specified. The species used in the alignment for each test are also shown. The log likelihoods for each model, *ω* estimates and p-values are also provided. See Methods for details.

**Supplementary Table 11.** Testing the direction of dependency between *PRDM9* and candidate genes *ZCWPW1, ZCWPW2, TEX15* and *FBX047*. Here, we asked whether we could reject a model of independent state transitions of *PRDM9* and a given candidate gene (e.g. *ZCWPW1*) in favor of a model in which state transitions of the candidate gene depend on those of *PRDM9* (model X). Next, we asked whether we could reject the null model in favor of a model in which the state transitions of *PRDM9* depend on those of the candidate gene (model Y). For comparison, we also provide results for the test shown in the main text, in which the alternative considered is that state transitions of *PRDM9* depend on those of the candidate gene and vice versa (also shown in Table 1). See Pagel, 1994 and the BayesTraitsV3.0.2 Manual for further description of these models and tests.

## Supplementary figures

**Figure S1.**
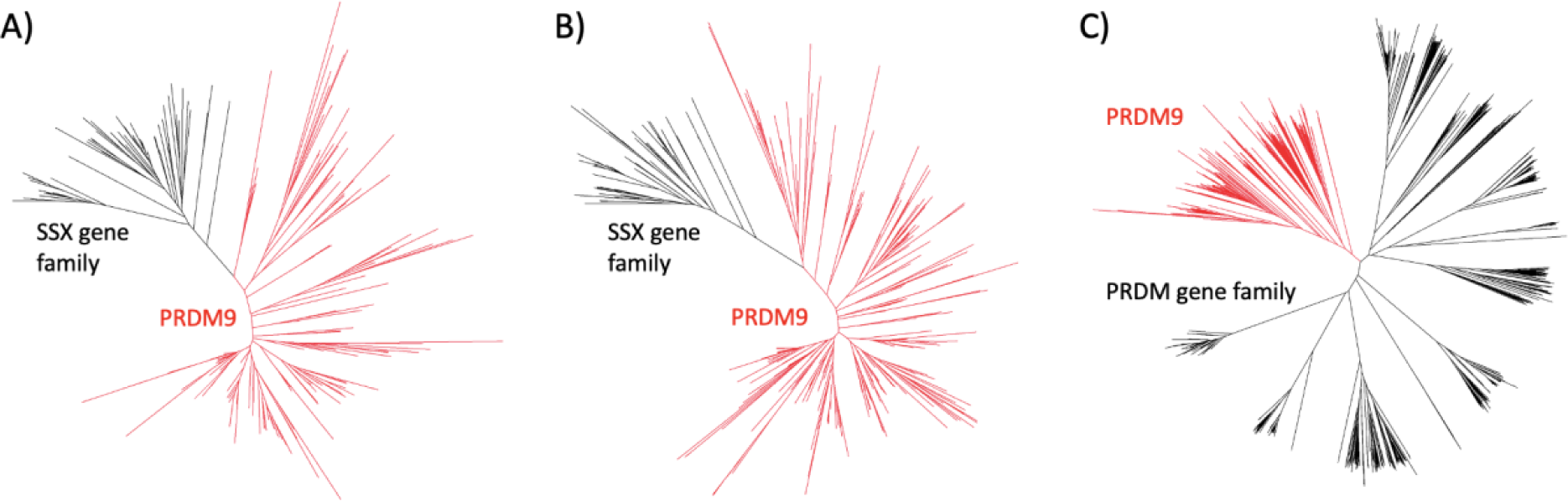
Guide trees created from our initial blastp search results for PRDM9 orthologs for **(A)** KRAB domains, **(B)** SSXRD domains and **(C)** SET domains. Genes were removed if they clustered with SSX genes in trees **(A)** or **(B)**, or if they clustered with PRDM gene family genes other than *PRDM9* or *PRDM7* in the tree **(C)**. Genes clustering with *PRDM9* and retained for subsequent analysis are shown in red.

**Figure S2.**
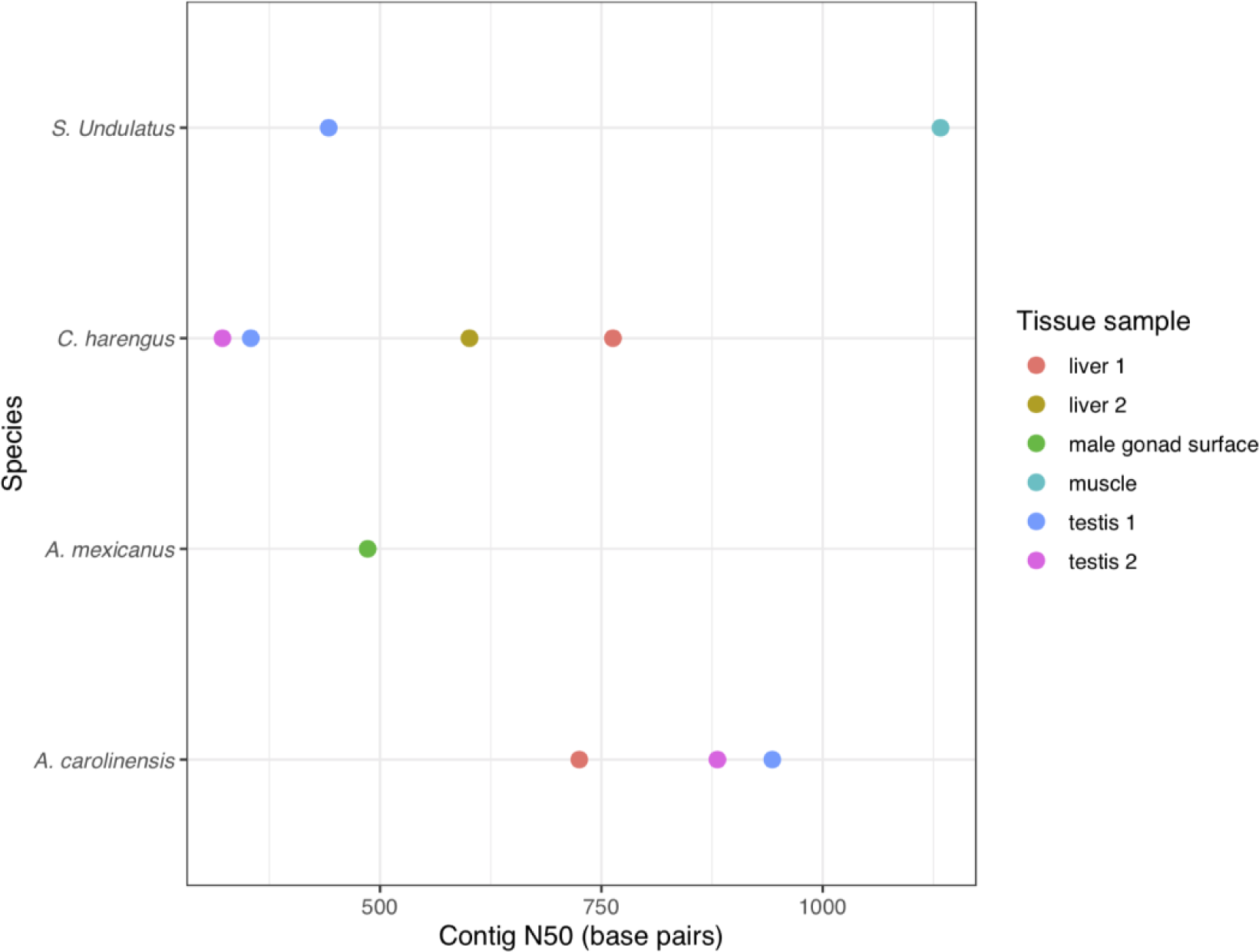
Contig N50 in base pairs as a statistic describing the quality of *de novo* transcriptome assemblies. Colors represent the different tissues used in the two lizard species (*S. undulatus* and *A. carolinensis*) and two fish species (*C. harengus* and *A. mexicanus*).

**Figure S3.**
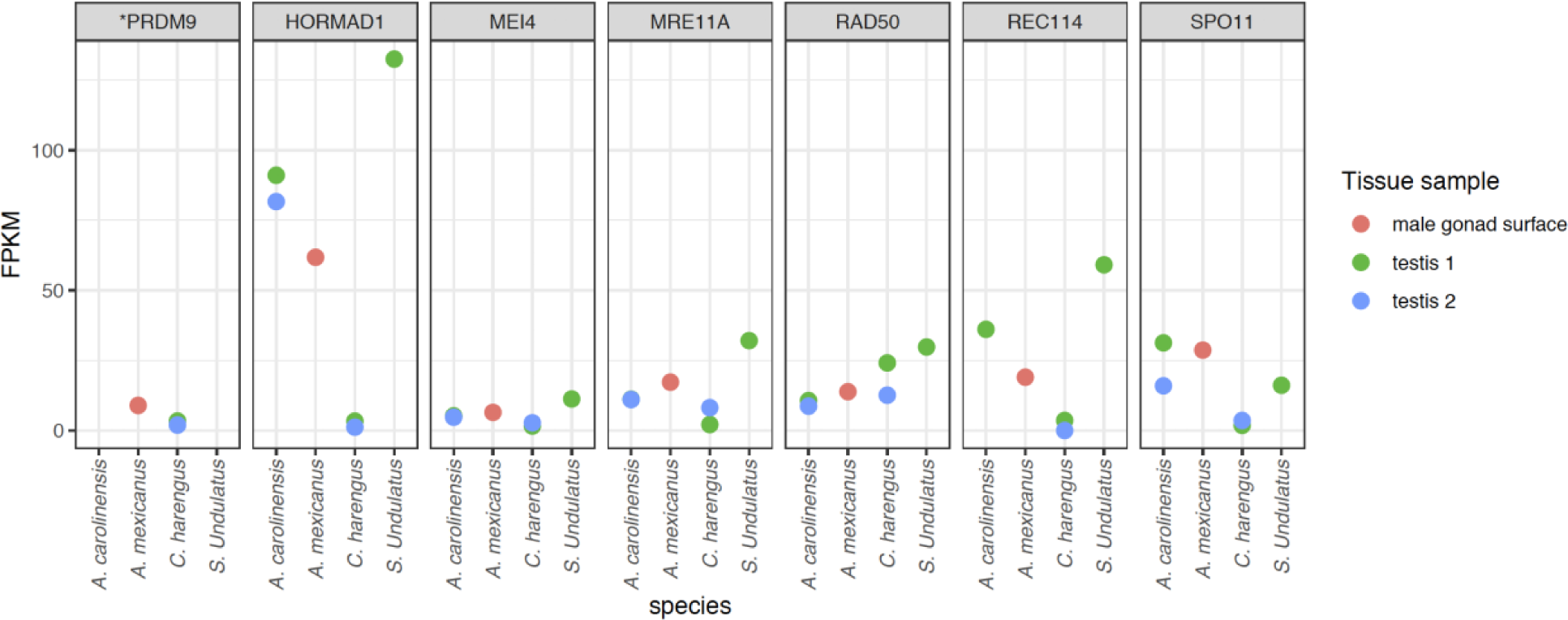
Expression levels of six core meiosis-related genes [32] across species and tissues. The y-axis corresponds to fragments per kilobase of transcript per million mapped reads (FPKM). Despite evidence for expression of the other six core meiotic genes, PRDM9 expression is not detected in *S. undulatus* and *A. carolinensis*.

**Figure S4.**
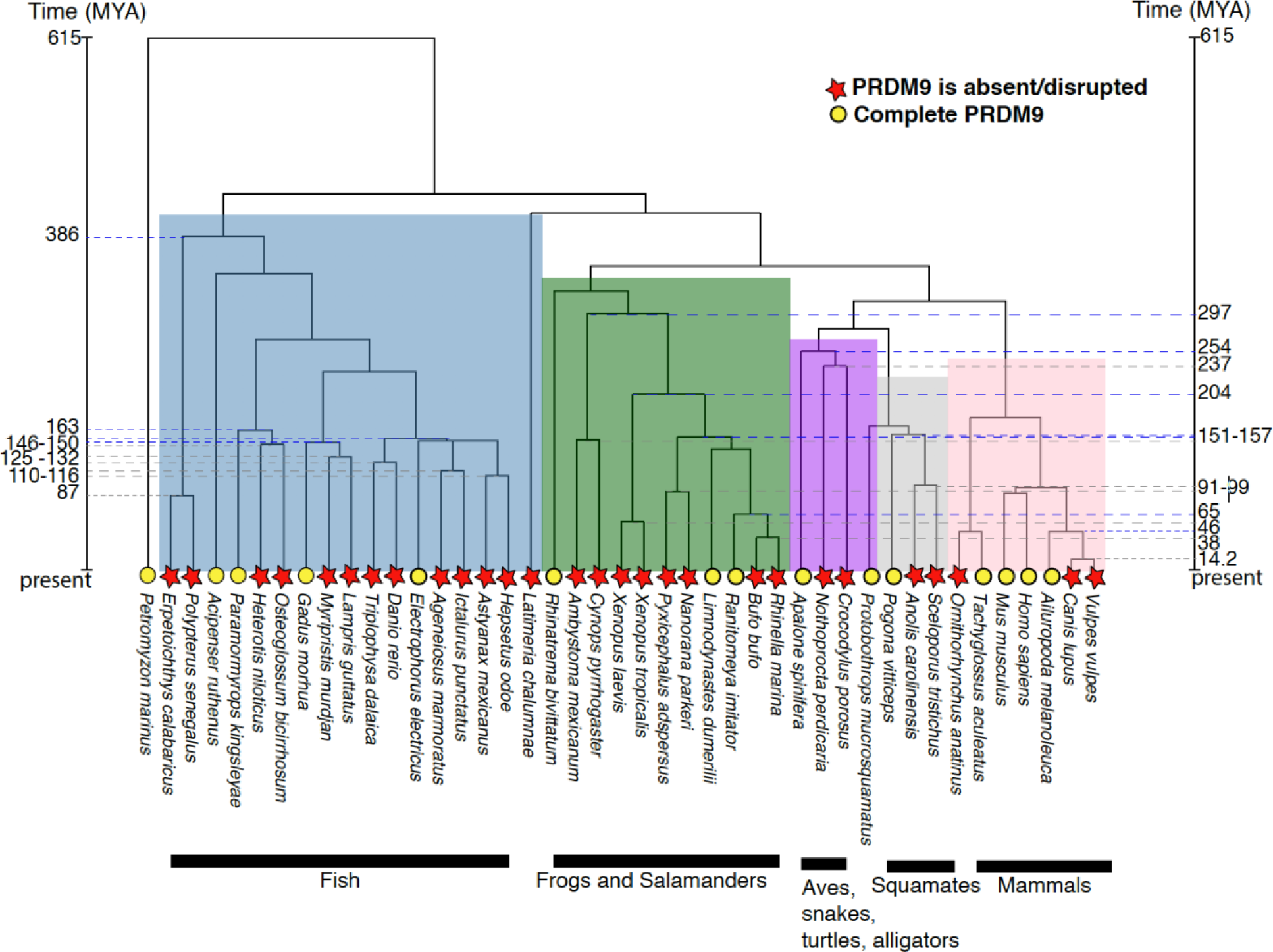
Phylogenetic distribution of *PRDM9* orthologs in vertebrates, using the phylogenetic tree and divergence dates obtained from Timetree [35]. A complete *PRDM9* was found in species marked with yellow circles. Species marked with a red star are ones for which we were unable to identify a complete PRDM9. The highlighted dates indicate the inferred timing of the multiple *PRDM9* losses. Dashed lines indicate the earliest (grey) and latest (blue) possible dates for each loss of *PRDM9*. The minimum date reflects the time to the most recent common ancestor amongst species without PRDM9, whereas the ‘maximum date’ is the time to the first common ancestor between species without *PRDM9* and the most closely related species with *PRDM9*. The most recent loss of *PRDM9* occurred either in the branch leading to canids, between 14.2 and 46 million years ago (Mya), or potentially the branch leading to platypus.

**Figure S5.**
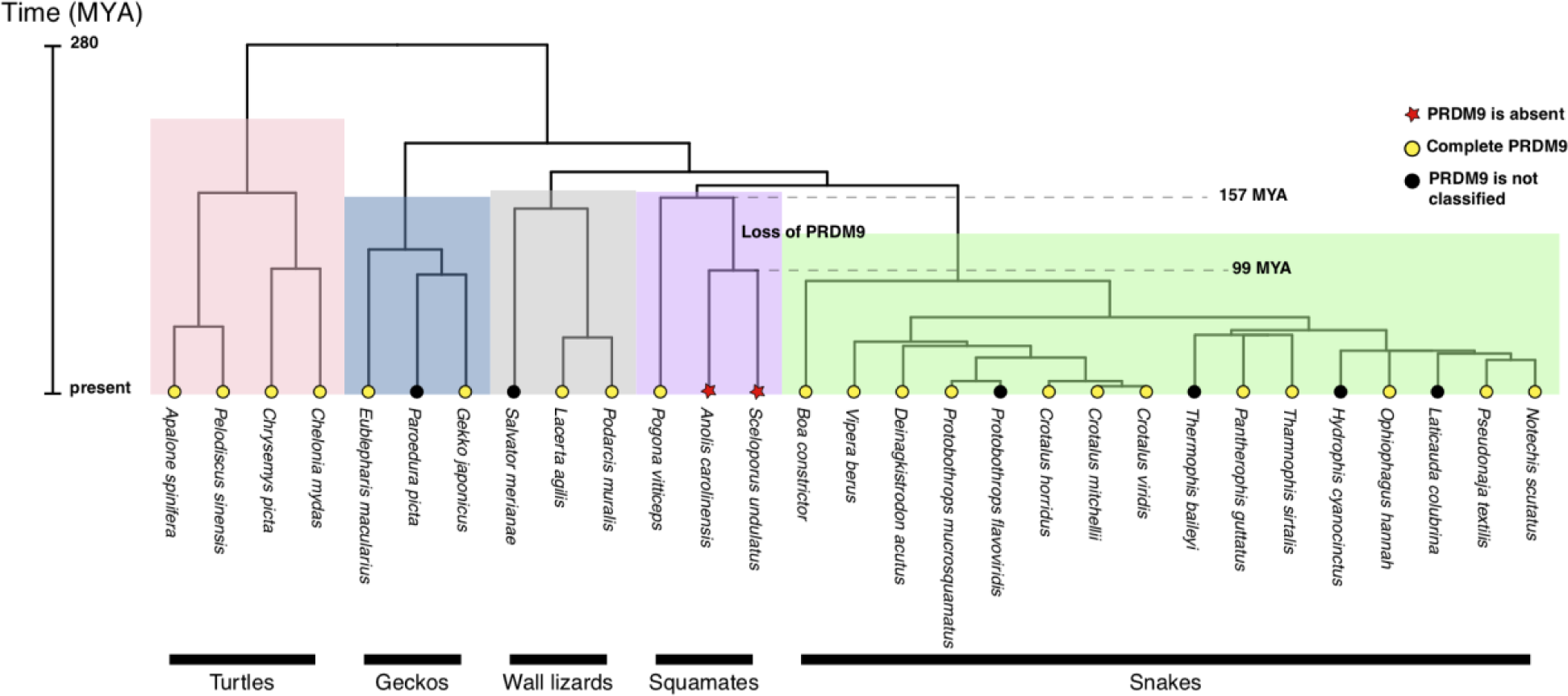
Phylogenetic distribution of *PRDM9* orthologs in reptiles, using the phylogenetic tree and divergence dates obtained from Timetree [35]. Species assigned with yellow circles carry a complete *PRDM9*. Species indicated with a red star are ones for which we were unable to identify PRDM9 expression in testis samples. Species indicated with a black circle are species for which Refseq is not available and *PRDM9* classification was therefore not conducted. Based on the phylogenetic relationship between *Anolis carolinensis* and *Sceloporus undulatus*, the *PRDM9* loss shared by these two species likely occurred between 99 and 157 million years ago (Mya).

**Figure S6.**
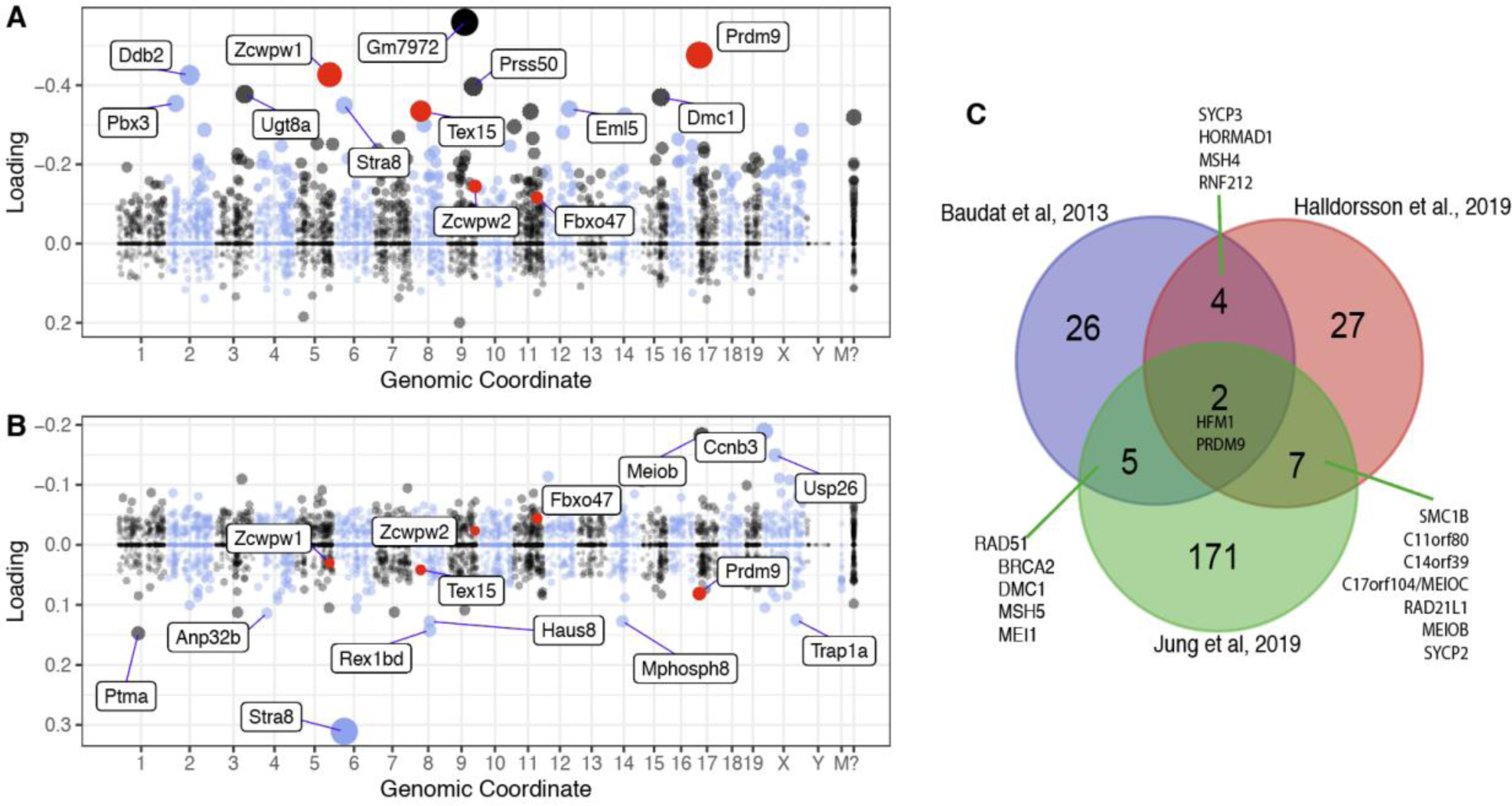
Meiosis-specific candidate genes. [31] inferred 46 principal components from single cell expression patterns during mouse spermatogenesis, which are thought to loosely correspond to regulatory programs. Shown in A-B are the two components in which PRDM9 is most highly expressed. The dot sizes are proportional to the square of the absolute value of the loading, so are indicative of higher expression within each component. PRDM9 and the five genes with p<0.05 in our phylogenetic analysis are shown in red. Mouse genomic coordinates are displayed. (A) Component 5 is the one in which PRDM9 has its highest loading; it is associated with double strand break formation and active during (pre)leptotene [31]. (**B**) Component 44 is the component in which PRDM9 has its second highest loading; this component is active during zygotene (Jung et al., 2019). (**C**) Intersection of candidate genes from three sources: (i) the top 1 percent of genes with highest loadings in component 5 (ii) genes associated with variation in recombination phenotypes in humans [33] and (iii) genes known to have a role in mammalian meiotic recombination from functional studies (as summarized in the review by [32]).

**Figure S7.**
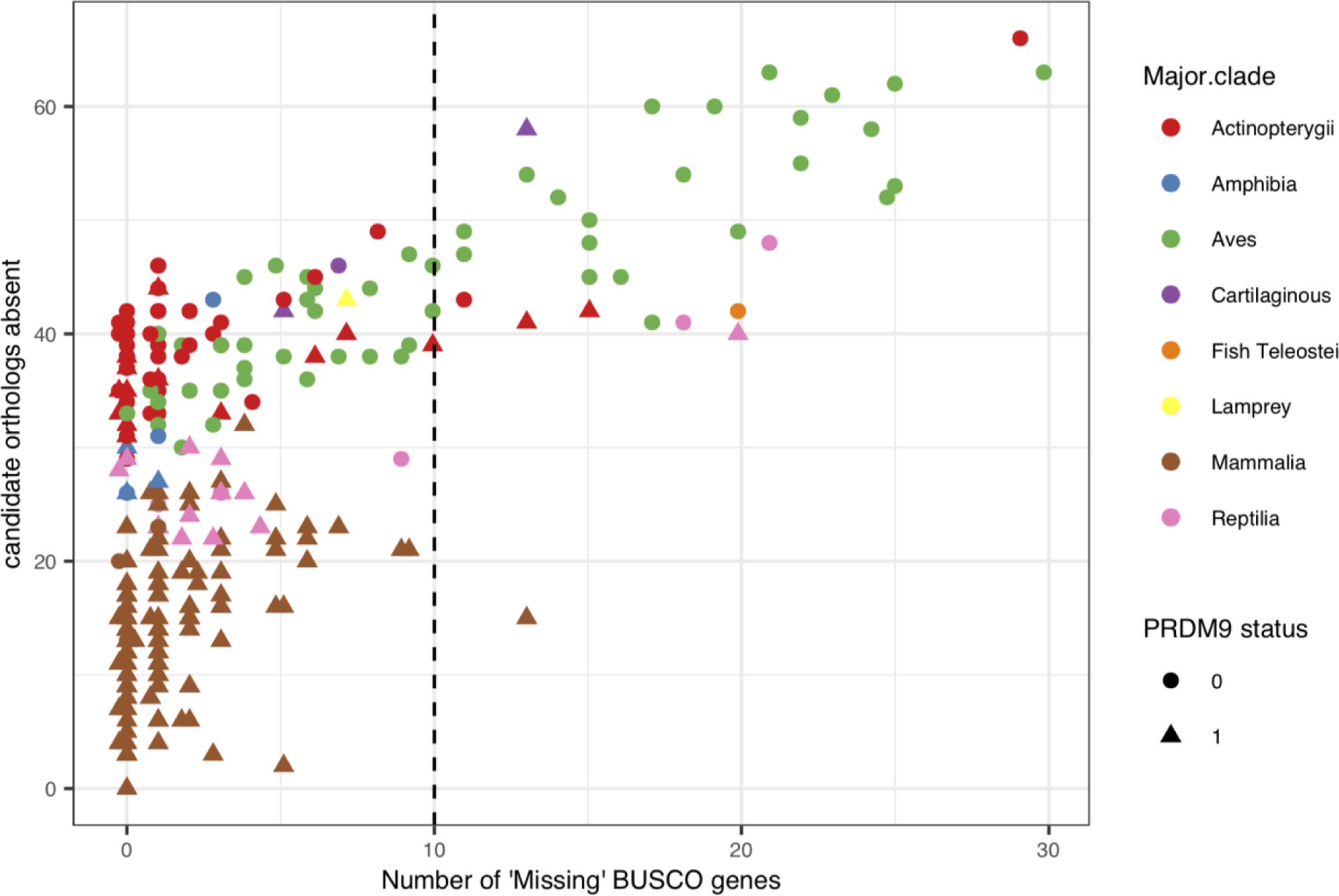
The relationship between the number of candidate genes that were absent in a genome assembly and the number of ‘Missing’ BUSCO genes [66] for that assembly, across species. BUSCO statistics were computed for the genomes of 339 species. The relationship is significant (Spearman’s rank correlation ρ = 0.5, p-value < 2.2e-16), suggesting that orthologs of candidate genes of interest might be missed in genomes with low BUSCO scores. In the phylogenetic tests, we therefore considered only species with 10 or fewer missing BUSCO genes (dashed line), leading 32 species to be excluded.

**Figure S8.**
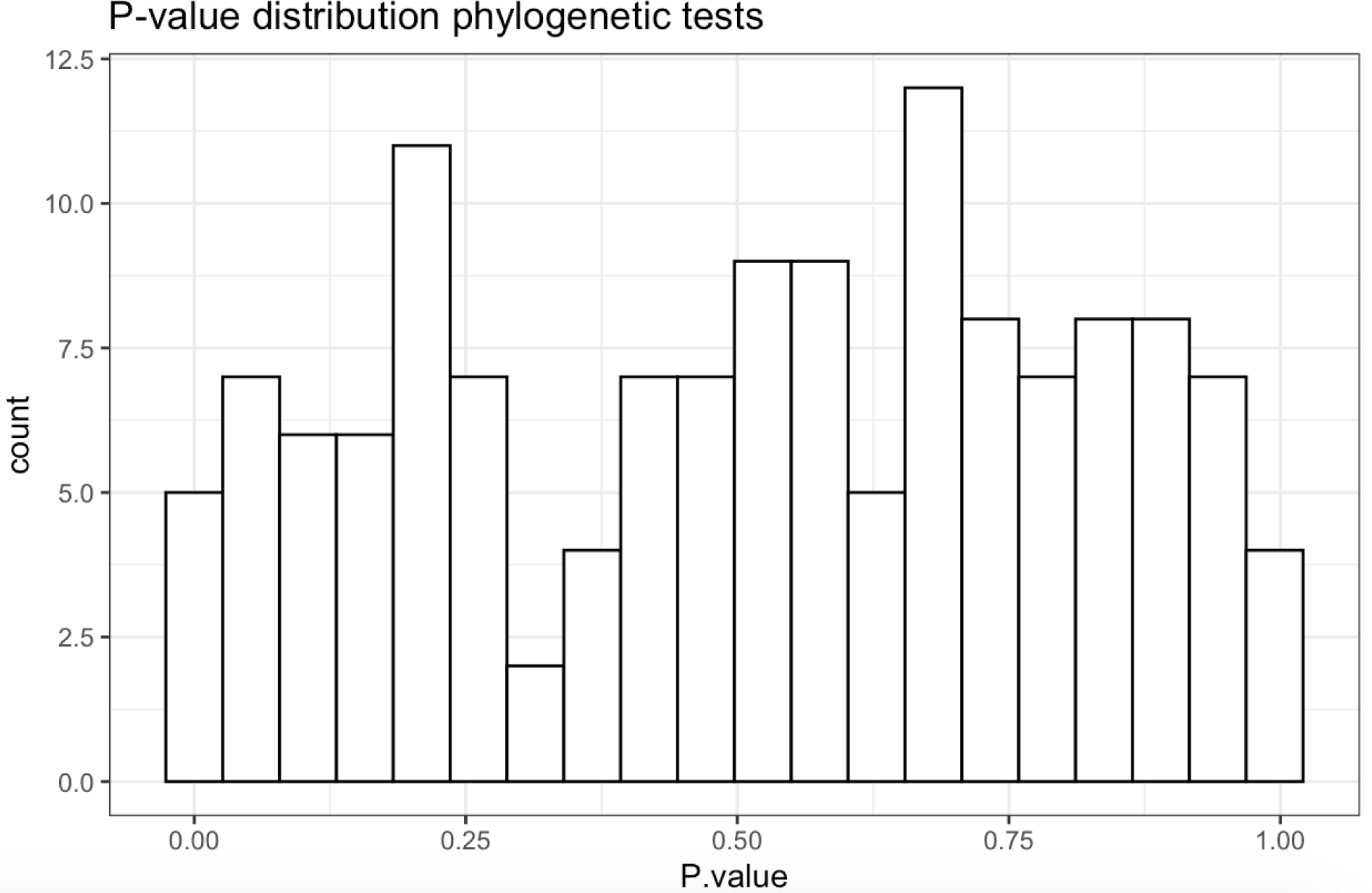
The distribution of p-values obtained across the 139 genes included in phylogenetic tests (individual p-values are available in Table S7).

**Figure S9.**
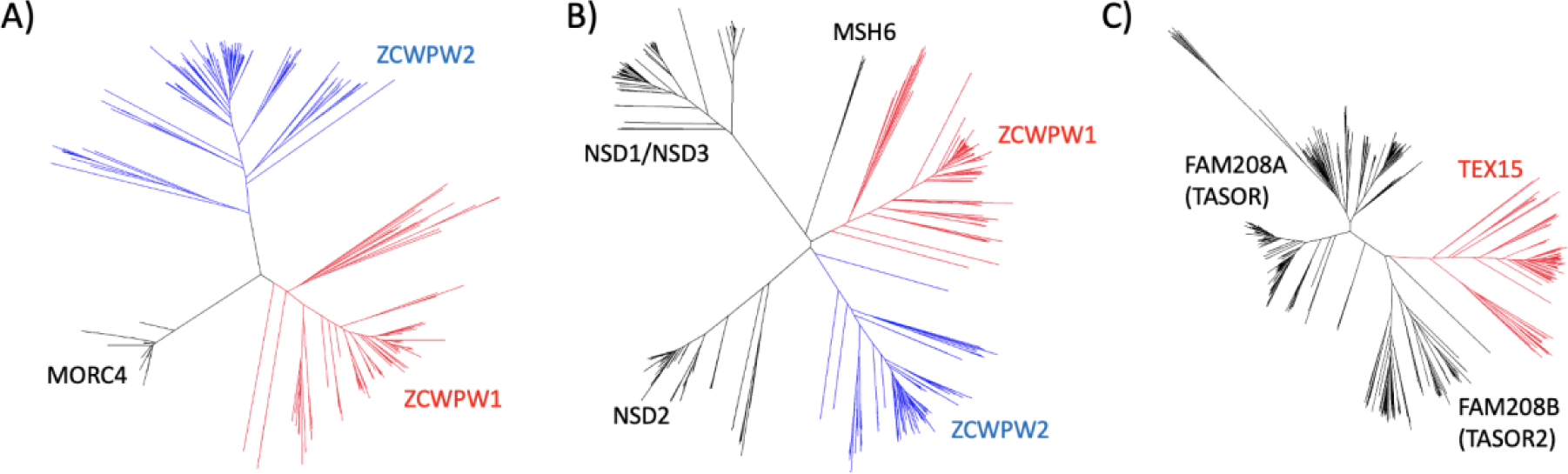
Guide trees created from our initial blastp search results for the zf-CW (**A**) and PWWP (B) domains of *ZCWPW1* and *ZCWPW2* orthologs, and the DUF3715 domain of *TEX15* orthologs. Genes were removed if they clustered with *MORC4* in tree **A**, *MSH6*, *NSD1*, *NSD2*, or *NSD3* in tree **B**, *FAM208A* or *FAM208B* in tree **C**. Genes clustering with *ZCWPW1*, *ZCWPW2* or *TEX15* and retained for subsequent analysis are shown in red or blue.

**Figure S10:**
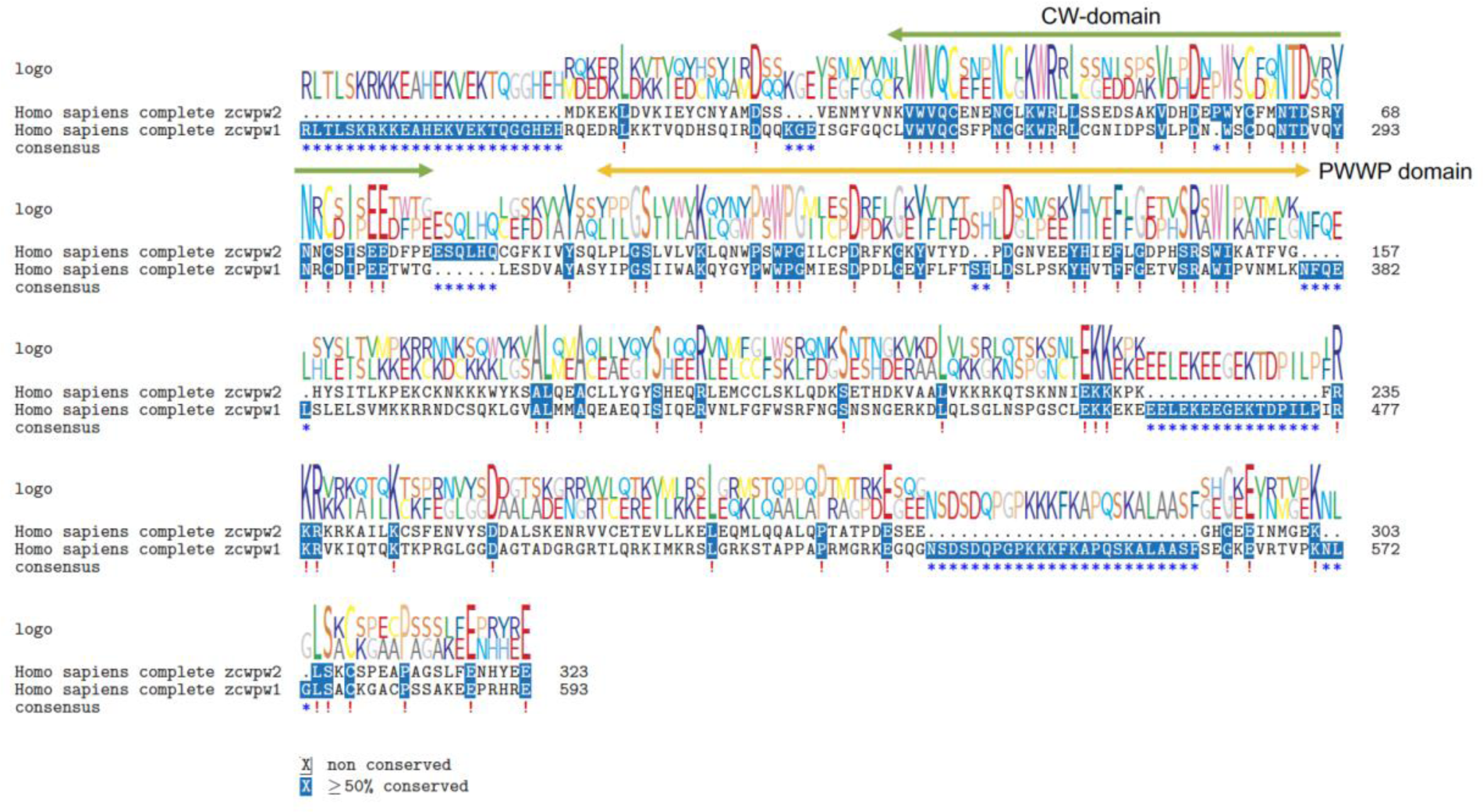
Amino acid sequence alignment between ZCWPW1 and ZCWPW2 proteins from humans. Superfamily domains are marked. In mice, the CW-domain (green arrow) recognizes different methylated states of lysine 4 on histone H3 (H3K4me) [77], while the PWWP domain (yellow arrow) recognizes methylated H3K36 histone tail [74]. The SET domain of PRDM9 tri-methylates both histones H3K4 and H3K36 [58].

